# Mechanisms Ensuring Fidelity of Family X DNA Polymerases in Programmed DNA rearrangements in Paramecium tetraurelia

**DOI:** 10.1101/2024.07.26.605286

**Authors:** Antonin Nourisson, Sophia Missoury, Ahmed Haouz, Marc Delarue

## Abstract

Repairing programmed DNA double-strand breaks (DSBs) is crucial in the lifecycle of *Paramecium tetraurelia*, especially during its sexual reproduction phase when its somatic highly polyploid macronucleus is lost. The formation of a new macronucleus involves Programmed Genome Rearrangements, introducing DNA DSBs at approximately 45,000 loci. *P. tetraurelia* employs a Non-Homologous End Joining (NHEJ)-related mechanism for the systematic repair of these DSBs. Four genes encoding DNA polymerases of family X are present in the genome, one of which was found recently to colocalize with other proteins of NHEJ. The question arises as to how they make almost no error. Here we show that these enzymes are most similar to metazoan DNA polymerase λ and exhibit high fidelity through two different molecular mechanisms. Using X-ray structure determination of polymerase lambda mutants recapitulating sequence determinants of *P. tetraurelia* PolXs, we find both a local conformational change that involves exchanging partners in a crucial salt bridge in the active site upon binding of correct dNTPs, and a larger conformational change involving the closure of Loop3. This stabilizes the template DNA in the active site, only in the presence of the correct incoming dNTP. Differences with human pol λ and pol β are discussed.

**GRAPHICAL ABSTRACT:** 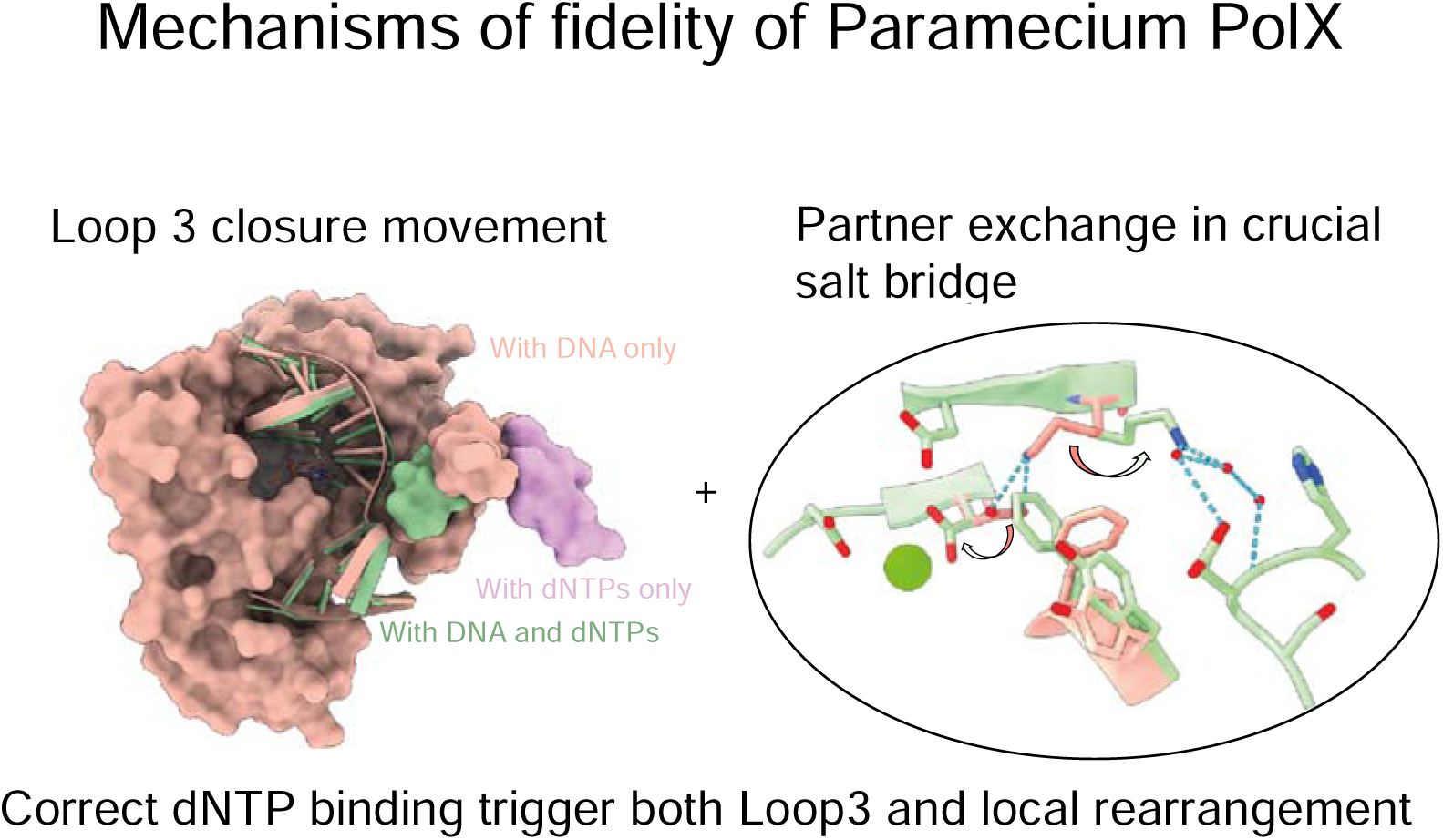

## INTRODUCTION

*Paramecium tetraurelia* exhibits nuclear dimorphism, with a single cell containing both a micronucleus (MIC), used for reproduction, and a macronucleus (MAC), utilized for gene expression (1). The MAC harbors up to 1600 copies of the 72 Mb somatic genome, while the diploid MIC genome (108 Mb) contains additional sequences, including Internal Eliminated Sequences (IES) (2). During the sexual reproduction cycle of *P. tetraurelia*, the MAC undergoes fragmentation, and a new MAC is formed from a copy of the MIC through extensive genome replication and rearrangements (3, 4). These large-scale alterations, known as programmed genome rearrangements (PGR), entail the precise removal of approximately 45,000 IESs. IESs are excised from the MAC DNA by a domesticated transposase known as PiggyMac (Pgm), which forms active complexes with non-catalytic Pgm-like proteins (5, 6). The DNA elimination complex induces double-strand breaks (DSB) specifically at dinucleotide 5’-TA-3’ sites flanking the IES, followed by the elimination of the 5’ terminal nucleotide (**Figure 1A)**. Recent studies revealed the participation of a specific Ku70/80 heterodimer in this complex (7) along with the XRCC4-Ligase IV complex (8–10), in a specialized NHEJ mechanism dealing with programmed DNA DSBs. While the NHEJ system is a priori capable to ensure fidelity in coordinated DNA DSB repair, especially in the presence of cohesive ends, (11, 12), the removal of the 5’ nucleotide at each DNA end following DSB induction necessitates an additional gap-filling step before ligation, involving at least one DNA polymerase to make the two DNA ends compatible (**Figure 1A)**. These findings prompt the question of the fidelity of the DNA polymerase involved (13, 14).

**Figure 1.**
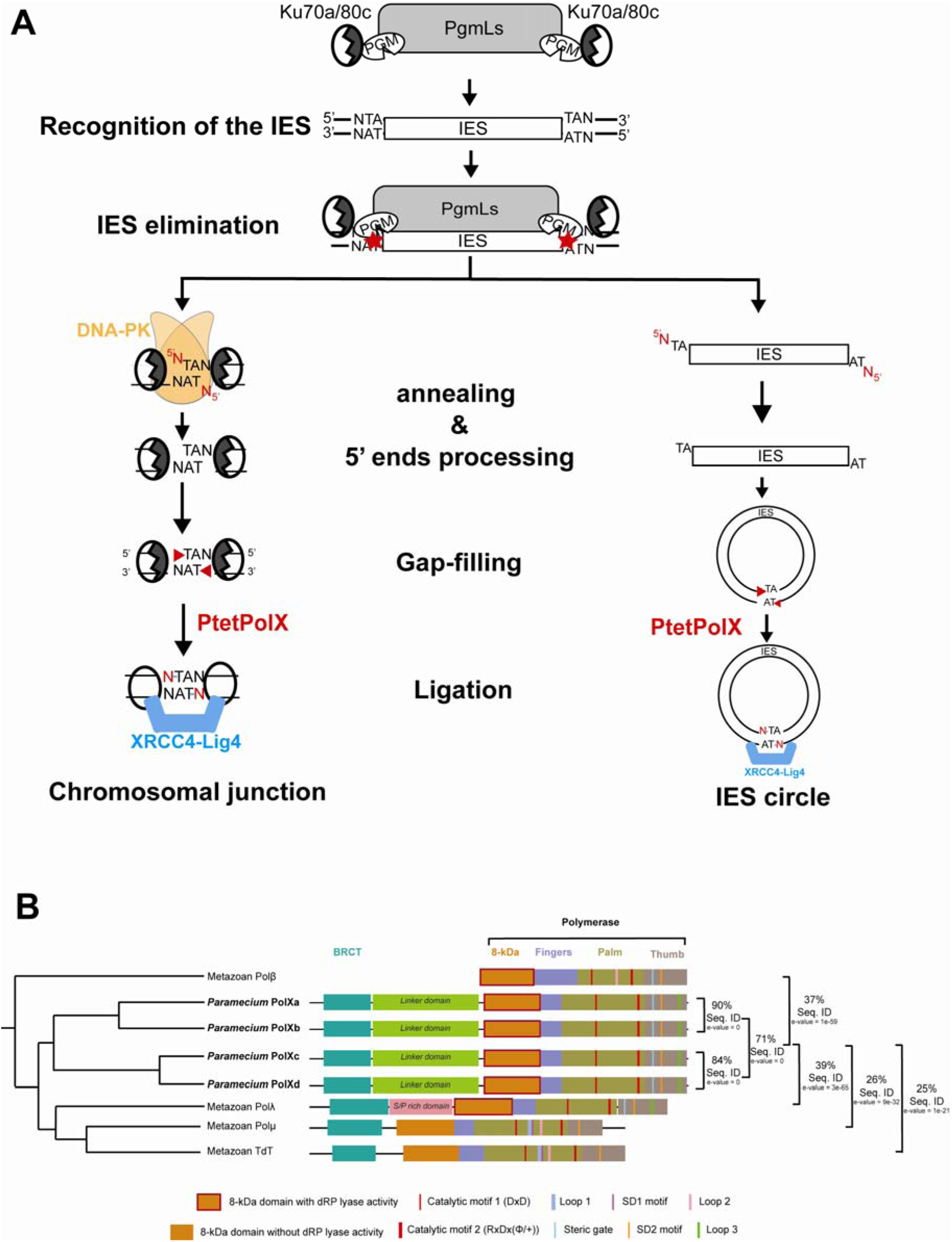
Role of PolXs in IES elimination during Programmed Genome Rearrangements in *P. tetraurelia*. A: After IES elimination by PiggyMac (PGM), the introduced Double Strand Breaks (DSB) are repaired by the NHEJ repair pathway, involving Ku70a/80c, DNA-PKcs, PolX and XRCC4-Ligase IV. *Paramecium* PolXs (Ptet-PolXs) are involved in the gap-filling step, during which their role is to accurately synthesize the missing base on each strand. B: *Paramecium* PolXs display a domain composition and sequence motifs similar to metazoan PolXs, especially the NHEJ-related Polλ and Polμ as well as V(D)J-related TdT. They share high sequence identity to one another and are divided in two groups (PolXab and PolXcd). Their closest homolog among metazoan PolXs is Polλ, but they are also homologs of Polβ, despite their additional BRCT domain. The domains are described under the figure, and the most important sequence motifs are indicated.

Four DNA PolXs genes, namely *POLXa, b, c*, and *d*, can be identified in the genome of *P. tetraurelia.* In addition, colocalization of one of them, polXa, with the other proteins of NHEJ (Ku and X4L4) has just been described (https://www.biorxiv.org/content/10.1101/2024.07.18.604053v1). Here we will focus on these four PolXs.

In metazoan (15), the gap-filling step is performed by DNA polymerases of family X (PolX) such as Polλ (16) or Polµ (17). Other members of the PolX family participate in short-range base excision repair (BER) (Polβ (18)), or a special version of NEHJ, namely V(D)J recombination (Terminal deoxynucleotidyl transferase (TdT) (19)). Both Polλ and Polµ are capable to deal with a wide spectrum of nucleic acids substrates resulting from different scenarii of DNA DSBs (20).

Apart from the N-terminal BRCT domain followed by a linker domain, PolX proteins consist of an 8-kDa domain and a DNA polymerase catalytic domain (**Figure 1B**). The 8-kDa domain aids in recognizing the 5’ phosphate DNA ends and exhibits a deoxyribose phosphate (dRP) lyase activity in Polβ and Polλ (21, 22). The DNA polymerase catalytic domain is composed of 3 subdomains (fingers, palm and thumb) and contains the catalytic aspartate triad, a steric gate motif (YFTGS, implicated in discriminating NTPs from dNTPs), three loops and two Sequence Determinant (SD1 and SD2) motifs (**Figure 1B**). Loop1 and (SD1) motif are specific to Polµ and TdT (23) ; their role is to help in the stabilization of a 5’ recessing template strand upstream the DSB, followed by a short microhomology (MH) region (24, 25), (26). The SD2 motif sequence is unique to each group of PolX. In TdT, the SD2 motif has been proposed to bind an additional divalent Zn^2+^ ion, but its precise role remains unclear (27). For TdT and Polμ point mutations in this SD2 motif deeply affect the polymerase activity (27, 28). In Polλ, the residues equivalent to Loop1 are involved in fidelity by controlling the placement of the primer DNA in the active site (29). The role of the SD2 motif has been shown to participate in the fidelity mechanism of Polβ (30). Loop3 is unique to Polλ and related homologues; its function is not currently known, but it has been observed by x-ray crystallography to undergo a large conformational change during substrate fixation and catalysis in the active site (31). In the presence of a correct incoming dNTP, Loop3 relocates closely to the DNA template strand and stabilizes the active site (31). Conversely, with incorrect dNTPs, Loop3 appears to be flexible, making it challenging to see it in crystal structures. In these instances, it does not interact closely with DNA, which is sub-optimally positioned in the active site for catalysis. Altogether, these recent kinetic crystallography experiments of polλ show that Loop3 likely plays a role in catalysis and potentially contributes to fidelity in Pol λ, but this requires further experimental functional evidence.

Notably, PolXs enzymes involved in NHEJ typically share a permanently closed conformation (32) and display low fidelity, particularly Polµ and TdT (33). On the other hand, the BER-associated Polβ displays the highest fidelity of metazoan PolX, which is mainly attributed to a large conformational change occurring when the right dNTP binds (34). In the presence of DNA and an incorrect incoming dNTP, the catalytic domain is in an open conformation with its thumb subdomain located away from DNA, and R258 forms a salt bridge with catalytic D192, diverting it from the active site and preventing catalysis (30). However, when DNA and the correct dNTP are present in the active site, the enzyme adopts a closed form, in which the thumb subdomain closes and stabilizes DNA in the active site, and the side chain of F272 (involved in the steric gate) now separates R258 and D192. In this active conformation, R258 forms a salt bridge with the glutamate residue of the SD2 motif (NEY), and D192 is now available for catalysis. (34). Interestingly, in Polβ the dNTP and DNA binding sites are not preformed, which in contrast with the apo form of Polμ and Polλ where there is a preformed DNA binding site, and with the apo form of Polλ that binds dNTPs in absence of DNA (35). The mutator effect of this prestabilization of dNTPs is counterbalanced by an hydrophobic cluster that regulates the transition to an active state of the catalytic site (35, 36).

The four DNA PolXs genes, *POLXa, b, c*, and *d* stemm from two whole genome duplications (WGD) (37). These genes are paralogues and exhibit varying degrees of sequence identity. At the protein sequence level, PolXa and PolXb share 90% sequence identity, while PolXc and PolXd share 84% of sequence identity, and the two subgroups collectively share 71% of sequence identity (**Figure 1B**). Notably, PolXa is overexpressed during PGR in *P. tetraurelia* (38). They share sequence similarities with both human Polβ (37% sequence identity) and Polλ (39% sequence identity), mainly restricted to the polymerase domain, and they exhibit common features in their domain organization, including a putative N-terminal BRCT domain, a linker domain, an 8-kDa domain containing residues possibly involved in dRP lyase activity, and a catalytic DNA polymerase domain resembling DNA polymerases λ and β (**Figure 1B**). The BRCT domains of *P. tetraurelia* PolXs do not align well with the ones of metazoan PolXs by BlastP searches but AlphaFold confidently predicts the presence of a BRCT domain in all four variants (**Suppl. Figure S1)**. However, the linker domain appears to be different from the one in Pol λ, as it is highly SP-rich in Pol λ and predicted by PONDR to be disordred (confirmed by AlphaFold), while it is not in *P. tetraurelia* PolXs and could be partially ordered according to AlphaFold (**Figure S1)**. The BRCT domain is responsible of the association of the PolX with the rest of the NHEJ complex, especially the Ku70/80 heterodimer. Here we ask what makes the polymerase domain of these DNA PolX special with respect to human pol μ and pol λ and how they could contribute to the high fidelity of *P. tetraurelia* ’s NHEJ process, ensuring fully reliable repair after the excision of IES by Pgm and their recruitment by Ligase IV (8, 12).

In the following we present the functional characterization of *Paramecium* PolXs (Ptet-PolX), and show that their enzymatic kinetics parameters are similar to Polλ’s, but with a better fidelity relying only on their catalytic domain. By structural studies of mutants of human pol lambda mutants carrying sequence determinants of *P. tetraurelia* polX identified by careful sequence analysis, we found two mechanisms that could explain the fidelity of the Ptet PolXs: (i) a local induced fit with an exchange of partner in a salt bridge in the catalytic site, like Polβ (but without its major domain rearrangement and open/closed transition), and (ii) the closing of Loop3, that could be described as a more global substrate-induced conformational change.

## MATERIAL AND METHODS

### Wild-type and mutant constructs of PolX

In this study, we purified different constructs of *Paramecium* PolXs. PolXd (ParameciumDB PTET.51.1.P1010039) was expressed and purified as a full-length construct (PolXdFL), while three constructs lacking residues 1-266 (including the N-terminal BRCT domain and the linker domain) were generated: PolXaΔNter (WT sequence: ParameciumDB PTET.51.1.P0210235), PolXbΔNter (WT sequence: *Paramecium*DB PTET.51.1.P0360066), and PolXdΔNter. Additionally, three mutant versions of PolXaΔNter were created: PolXaΔNter-K534A (mutation K534A), PolXaΔNter-K534R (mutation K534R), and PtetLoop3β where residues 581 to 588 (Loop3) were replaced by the corresponding sequence of Pol β (GVA).

Wild-type versions of human Polλ (Uniprot Q9HBN3) and Polβ (Uniprot P06746) were also produced. Furthermore, five mutant versions of human Polλ were generated, all featuring the following mutations originally described in Jamsen et al. (32): Δ1-241 (ΔNter), replacement of residues 464-472 (Loop1) with the equivalent sequence in polβ (KGET), and C544A. This construct will be referred to as λref in the following. The specific mutations in the other constructs were as follows (**Table 1)**: I492R (λmutR), I492R and residues 528 to 530 (SD2 region) replaced by the corresponding –much shorter-polβ sequence NEY (λSD2β), a single mutation I492K (λmutK), and two mutations - I492K and E529D (λSD2Ptet). Additionally, another construct based on human Polλ, named λloop3β, was produced and purified, with residues 539-547 (Loop3) replaced by the corresponding sequence of Polβ (GVA).

**Table 1.**
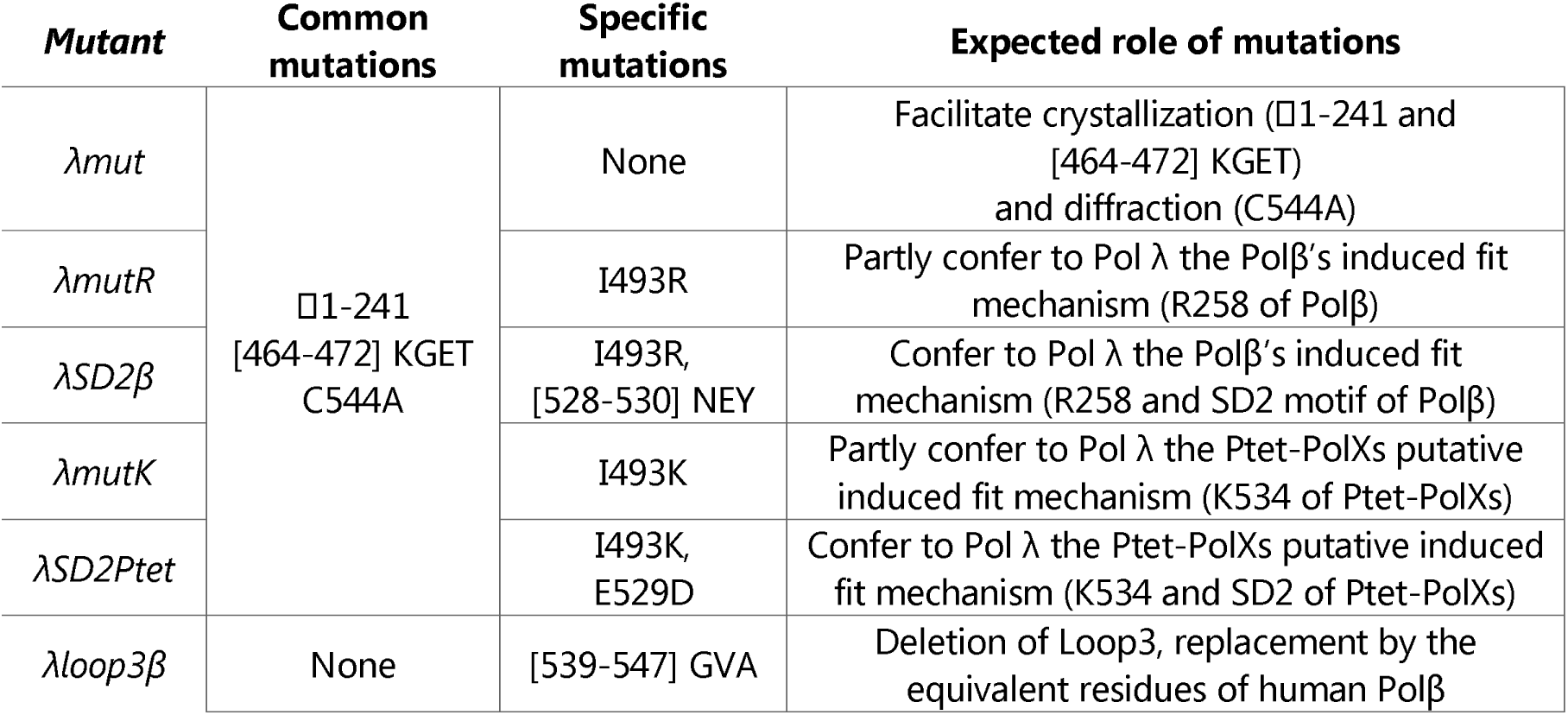
Mutant constructs of human Polλ used for enzymatic and structural studies of the fidelity mechanisms of Ptet-PolXs.

All the codon-optimized sequences for WT constructs are indicated in **Supplementary Table S1**, and mutated constructs and used primers are described in **Supplementary Table S2**.

### Cloning, overexpression, and purification of Paramecium PolXs (Ptet-PolXs)

The synthetic genes encoding Ptet-PolXs were optimized for expression in *E. coli* and synthesized using ThermoFisher’s GeneArt service. These genes were then cloned into a modified pRSF1-Duet expression vector with a cleavable N-terminal 14-histidine tag. For the PolXdFL and PolXbΔNter constructs, cloning was done using New England Biolabs and Anza (Thermo Fisher Scientific) restriction enzymes, while PCR and NEBuilder HiFi DNA Assembly (New England Biolabs) were utilized for the other constructs. The mutant constructs of PolXa□Nter were generated via site-directed mutagenesis performed by PCR (primers are indicated in **Supplementary Table S2)** and using the KLD enzyme kit (New England Biolabs).

*E. coli* BL21 Star (DE3) cells (Invitrogen) were transformed with the engineered plasmids and cultured at 37°C in LB medium with kanamycin resistance selection. Induction was carried out at OD = 0.6–1.0 with 1 mM IPTG, followed by overnight incubation at 20°C. After harvesting, cells were homogenized in buffer A (50 mM Tris-HCl pH 8, 600 mM NaCl, 10 mM imidazole), sonicated, and centrifuged to remove bacterial debris. The resulting lysate supernatants were treated with Benzonase (Sigma-Aldrich) and protease inhibitors (Thermo Fisher Scientific) before purification.

The proteins of interest were isolated by purifying the clarified cell lysates on a HisTrap column, using buffer A (50 mM Tris-HCl pH 8, 600 mM NaCl, 10 mM imidazole) as the washing buffer and 500 mM imidazole in the elution buffer. For the PolXbΔNter construct, a different washing buffer (25 mM Sodium Phosphate pH 8, 1 M NaCl) was utilized to avoid nucleic acid contamination. The collected proteins were then dialyzed to 75 mM NaCl and repurified on a HiTrap Heparin column with an elution at 1 M NaCl. Both purification columns were from Cytiva.

Protein purity was analysed using SDS-PAGE 4-15% or 4-12% gels with a molecular weight ladder (Precision Plus Protein, Biorad) as a control. The enzymes were concentrated using Amicon Ultra 30k MWCO centrifugal filters (Merck), flash frozen in liquid nitrogen, and stored directly at −80 °C until further use.

### Cloning, overexpression, and purification of human Polβ

The gene encoding human Polβ was commercially synthesized by Genscript and inserted into the modified pRSF1-Duet expression vector with a cleavable N-terminal 14-histidine tag. Production and purification followed the same protocol as described for the previous constructs, with the use of specific buffers: Buffer A (50 mM Tris-HCl pH 8, 500 mM NaCl, 10 mM imidazole), Buffer B (50 mM Tris-HCl pH 8, 500 mM NaCl, 500 mM imidazole), Dilution buffer (50 mM Tris-HCl pH 8), Buffer C (50 mM Tris-HCl pH 8, 100 mM NaCl), Buffer D (50 mM Tris-HCl pH 8, 1 M NaCl).

### Cloning, overexpression, and purification of human Polλ and mutant constructs

The gene expressing human Polλ was commercially synthesized by Genscript and inserted into the modified pRSF1-Duet expression vector with a cleavable N-terminal 14-histidine tag. Production and purification followed the same protocol as described for the previous constructs, utilizing the following buffers : Buffer A (50 mM Tris-HCl pH 8, 500 mM NaCl, 10 mM imidazole, 1 mM EDTA, 1 mM DTT, 5% glycerol), Buffer B (50 mM Tris-HCl pH 8, 500 mM NaCl, 500 mM imidazole, 1 mM EDTA, 1 mM DTT, 5% glycerol), Dilution buffer (50 mM Tris-HCl pH 8, 1 mM EDTA, 1 mM DTT, 5% glycerol), Buffer C (50 mM Tris-HCl pH 8, 100 mM NaCl, 1 mM EDTA, 1 mM DTT, 5% glycerol), Buffer D (50 mM Tris-HCl pH 8, 1 M NaCl, 1 mM EDTA, 1 mM DTT, 5% glycerol).

All mutant constructs were obtained by PCR (primers are indicated in **Supplementary Table S2)** and using the KLD enzyme mix (New England Biolabs) from the plasmid used for the expression of Hs Polλ, and purified similarly to the WT protein. An additional step of size-exclusion chromatography was performed on a HiLoad Superdex 200 16/60 PG gel filtration column (Cytiva) in Storage Buffer (50 mM Tris-HCl pH 8, 100 mM NaCl). All purified proteins were concentrated to 16 to 20 mg/ml and stored at −80 °C after flash freezing in liquid nitrogen until further use.

### Gap-filling and DSB-cis polymerase assays

Primer extension activity assays were conducted in a reaction buffer consisting of 50 mM Tris-HCl pH 7.5, 50 mM KCl, 5 mM MgCl_2_, 1 mM DTT, and 5% glycerol. Reaction solutions comprised 1 μM of templating oligonucleotide, 1 μM of FAM 5’-labelled DNA primer, and either 250 μM dGTP or a mixture of all four dNTPs (250 µM each).

For the gap-filling assay, the used oligonucleotides were the following: 5’-FAM-AATCACCAGTACGCCGTTGCGT-3’,5’-p-TATCGCCATGACGCGGTTCTGGTCC-3’, 5’-GGACCAGAACCGCGTCATGGCGATACACGCAACGGCGTACTGGTGATT-3’. Reactions were incubated for 30 minutes at 27°C with 1 nM of polymerase (Ptet-PolX) or for 5 minutes at 37°C with 50 nM of polymerase (HsPolλ). In the DSB cis assay, the oligonucleotides were the following: 5’-FAM-AATCACCAGTACGCCGTTGCGT-3’, 5’-p-TATCGCCATGACGCGGTTCTGGTCC-3’, 5’-TACACGCAACGGCGTACTGGTGATT-3’, 5’-GGACCAGAACCGCGTCATGGCG-3’. Reactions were incubated for 30 minutes at 27°C (the optimal growth temperature of *P. tetraurelia)* with various concentrations of PolXaΔNter. Prior to adding the protein, DNA was hybridized by heating up to 90°C and gradually cooled down to room temperature. For the primer extension assay, the used oligonucleotides were the following: 5’-FAM-AATTGTCATAAGCTTATGCG-3’, 5’-p-TATCGCCATGACGCGGTTCTGGTCC-3’, 5’-GGGGTAGCTGCGCATAAGCTTATGACAATT-3’. Reactions were incubated for 30 minutes at 27°C with 1 µM of polymerase (Ptet-PolX) or at 37°C with 50 nM of polymerase (HsPolλ).

Reactions were terminated by adding two volumes of a buffer containing 10 mM EDTA, 98% formamide, and 1 mg/mL bromophenol blue, and stored at -20°C. Products were preheated at 95°C for 10 minutes, separated using denaturing urea-polyacrylamide (18%) gel electrophoresis, and visualized by FAM fluorescence on a Typhoon FLA 9000 imager. All oligonucleotides were obtained from Eurogentec, Eurofins, or Biomers, dNTPs from Fermentas (Thermo Fisher Scientific), and chemicals from Sigma-Aldrich.

### dRP lyase assay

For the dRP lyase assay, DNA was prepared using the following procedure: a 31-mer DNA strand (5’-GTACCCGGGGATCCGTACAGCGCATCAGCTGCAG-3’) and its complementary U-containing DNA strand (5’-CTGCAGCTGATGCGCUGTACGGATCCCCGGGTAC-3’), each at a concentration of 50 μM, were hybridized as described above. Next, 4 picomoles of hybridized DNA were mixed with USER3 mix (New England Biolabs) in ThermoPol buffer (20 mM Tris-HCl pH 8.8, 10 mM (NH_4_)_2_SO_4_, 10 mM KCl, 2 mM MgSO_4_, 0.1% Triton® X-100), and the mixture was incubated for 2 hours at 65°C.

Subsequently, in a reaction volume of 10 μL, 1 μM of USER3-treated DNA was mixed with a 250 μM dNTP mix, 400 units of T4 DNA ligase, and 50 nM of each DNA polymerase (HsPolβ or PolXaΔNter), in an activity buffer (50 mM Tris-HCl pH 7.5, 5 mM MgCl_2_, 50 mM KCl, 1 mM DTT, 1 mM ATP). The mixture was then incubated for 30 minutes at 27°C (for Ptet-PolX) or 37°C (for HsPolβ) and analyzed by urea-PAGE in denaturing conditions as described previously.

### Enzymatic steady state characterization

DNA polymerization assays were conducted between 3 to 12 times using the methodology described above. Each assay utilized 1 μM of gap-filling DNA and 5 nM of DNA polymerases, with varying concentrations of dNTPs, and proceeded for 10 minutes at 27°C.

The resulting gels were subjected to analysis using ImageJ software, and quantification of the product DNA was carried out using Microsoft Excel. Enzyme velocity (measured in nM/min) was plotted against the concentration of dGTP. These plotted data points were then fitted to a nonlinear regression curve utilizing the Michaelis-Menten equation with GraphPad Prism 10 software. From the fitted curves, values for k_obs_ and K_M_ were obtained.

### One-point single turnover fidelity assay

The fidelity assays were conducted following the protocol established by Fiala *et al.* (39) in a buffer comprising 50 mM Tris-HCl (pH 8.4), 5 mM MgCl2, 100 mM NaCl, 0.1 mM EDTA, 5 mM DTT, 10% glycerol, and 0.1 mg/mL BSA. Each reaction mixture contained 30 nM of gap-filling DNA, 120 mM of each dNTP, and 120 nM of DNA polymerases. The reactions were carried out at 37°C (for HsPolλ) or 27°C (for Ptet-PolX) for 50 minutes. Following completion, the reactions were stopped and subsequently analyzed as described previously.

### Clustering of the PolX family

6500 PolXs sequences were obtained by PSI-BLAST (40) of the NCBI (41) non redundant database, using the sequence of Ptet-PolXa as a probe. Sequences were sorted by query coverage, and the last chosen sequence displayed the following parameters: query coverage = 52%; e-value = 2.10 ; % identity = 32,02%. To obtain representative sequences for the bacterial PolXs and yeast PolIV groups, three other PSI-BLAST were performed, using as probes PolX from Thermus thermophilus [NCBI ID: WP_096410530.1], PolX from Deinococcus radiodurans [NCBI ID: WP_010887112.1] and PolIV from Saccharomyces cerevisiae [NCBI ID: AJP37443.1]. From each search, the top 250 sequences were chosen.

The FASTA file containing 7250 final sequences was processed by the CLANS web-utility from the MPI Bioinformatics Toolkit (42, 43). The clustering simulation was conducted using the Java version of CLANS (44) with default parameters. The sequences were randomly distributed in 3D space, converging to individual clusters after several hundred steps of the simulation. The simulation was let to run for a total of 20,000 steps, during which no further modification of the positions appeared. The simulation was run without applying a p-value cut-off: additional simulations with cut-offs (10^−10^ and 10^−20^) resulted in a shrinking and fragmentation of the clusters, leading to the impossibility to observe known relationships between PolX. Clusters were curated manually. Several independent simulations converged to almost identical cluster distributions, with no discrepancy regarding the critical details. Simulations not enriched with bacterial or yeast PolXs sequences resulted in equivalent distributions.

### Sequence multialignment

A multialignment of various eukaryotic PolXs sequences was computed using PSI-Coffee (45) and encompass all regions of interest located in the DNA polymerase domain. Graphical representation of the multiple sequence alignments was generated using ESPript 3 (46), with secondary structures from the human Polλ (PDB 7M43) provided for reference.

### Domain prediction

The on-line version of PONDR (47) was used to predict potential disordered regions in *P. tetraurelia* and human PolXs. AlphaFold3 (48) was used to predict their structure. The PAGSTscore was calculated over a sliding window of 10 residues and smoothed over a window of 5 residues.

### Crystallization

For crystallization of λRef and its variants, the DNA substrate was prepared in the following way: an 11-mer template oligonucleotide (5′-CGGCAGTACTG−3′) was annealed with a 6-mer upstream primer oligonucleotide (5′-CAGTAC-3′) and a 5′-phosphorylated downstream 4-mer primer oligonucleotide (5′-pGCCG-3′) in a 1:1:1 ratio. The crystallization plates were prepared according to the procedure outlined by Jamsen *et al.* (31). In brief, the protein (16 mg/mL) was combined with DNA in a 1:2 molar ratio and incubated at 4□°C for 2□h. Following the addition of 2□mM dNTP, either dTTP (matched) or dCTP (unmatched), and 10□mM CaCl_2_, the mixture was further incubated for 2□h on ice. Crystallization drops were set up at 4°C by mixing in a 1:1 ratio the Polλ-DNA-dNTP ternary complexes with a crystallization solution consisting of 20 mM Bicine pH 7.5, 300 mM Na-K tartrate, and 14-20% PEG 600/1000/10K/20K/Smear Low/Smear Medium/Smear High. For the crystallization of the mutant λmutK, an optimization of the best condition (20 mM Bicine pH 7.5, 300 mM Na-K tartrate, 17.5% PEG 20K) was conducted by screening additives (using the Additive Screen HT kit #HR2-428, Hampton Research) and lowering the protein concentration to 10 mg/mL. The resulting crystals were flash-frozen in liquid nitrogen after being cryo-protected with the crystallization solution supplemented with 25% ethylene glycol.

### Data collection and refinement

X-ray crystallographic data were collected at the SOLEIL synchrotron (Saint-Aubin, France) utilizing beamlines PX1 and PX2A. The data were processed using XDS (49) with the XDSME pipeline or autoPROC (50). The strong diffraction anisotropy of λSD2β dataset was taken into account with StarANISO (Staraniso GlobalPhasing, https://staraniso.globalphasing.org). Subsequently, the structures were solved by PHASER (https://www.globalphasing.com/) molecular replacement using a HsPolλ model (PDB ID: 7M43) as template and refined in Buster (51), with manual reconstruction performed using Coot (52). Details for all datasets are summarized in Table 2.

**Table 2.** Crystallographic statistics for datasets of crystals of pol lambda mutants with dNTP insertion site occupied opposite A in the presence of Ca^2+^. Data in the highest resolution shell is shown in the parenthesis. *Values calculated after truncation by STARANISO. Estimated resolution limits along the three crystallographic directions of the reciprocal lattice a*, b*, c*. ** Values obtained with MolProbity.

| | λmutR (dTTP) | λSD2β | λmutK d(TTP) | λSD2Ptet (dTTP) | λSD2Ptet (dCTP) |
| --- | --- | --- | --- | --- | --- |
| Data collection |  |  |  |  |  |
| Space group | P 21 21 21 | H32 | P 21 21 21 | P 21 21 21 | P 21 21 21 |
| a, b, c (Å) | 56.28 62.51 140.22 | 149.904 149.904 272.154 | 56.03 62.49 141.37 | 56.4 62.47 139.57 | 56.35 62.76 139.72 |
| $\alpha$, $\beta$, $\gamma$ (°) | 90 90 90 | 90 90 120 | 90 90 90 | 90 90 90 | 90 90 90 |
| Wavelength (Å) | 0.9801 | 0.9801 | 0.9801 | 0.9801 | 0.9801 |
| Resolution (Å) | 46.74 - 2.32 (2.38 - 2.32) | 117.173 - 3.543 (4.018 - 3.543) | 24.04 - 1.916 (2.118 - 1.916) | 46.55 - 2.12 (2.18 - 2.12) | 46.69 - 2.737 (2.987 - 2.737) |
| Estimated resolution limit (Å)* |  | 5.491 5.491 3.344 |  |  |  |
| R<sub>pim</sub> | 0.040 (0.629) | 1.514 (3.179) | 0.052 (0.807) | 0.038 (0.69) | 0.204 (0.845) |
| Completeness (%) | 99.9 (98.8) | 100 (100) | 93.3 (63.9) | 99.8 (97.5) | 86.4 (48.5) |
| Multiplicity | 13.3 (13.7) | 18.2 (19.2) | 11.9 (11.7) | 13.4 (13.7) | 9.3 (9.2) |
| I/$\sigma$(I) | 17.85 (1.99) | 5.00 (0.70) | 11.0 (1.70) | 16.6 (1.6) | 4.6 (1.40) |
| CC1/2 | 99.9% (82.0%) | 65.7 % (63.9%) | 99.6% (50.4%) | 99.9% (79.4%) | 97.5% (54.8%) |
| R<sub>pim</sub>* |  | 0.658 (1.192) |  |  |  |
| Completeness (%)* |  | 91.7 (74.2) |  |  |  |
| Multiplicity* |  | 17.7 (15.2) |  |  |  |
| I/$\sigma$(I)* | | 7.80 (1.80) | | | |
| CC1/2* |  | 80.5% (74.7%) |  |  |  |
| Refinement |  |  |  |  |  |
| No. of reflections | 22111 | 5991 | 25444 | 28561 | 8572 |
| R<sub>work</sub> / R<sub>free</sub> | 0.208 / 0.256 | 0.263 / 0.292 | 0.204 / 0.245 | 0.193 / 0.224 | 0.209 / 0.259 |
| No. non H atoms |  |  |  |  |  |
| Macromolecules | 2459 | 5036 | 2312 | 2514 | 2497 |
| Ligands | 47 | 0 | 31 | 47 | 28 |
| Solvent | 303 | 21 | 278 | 306 | 76 |
| Protein geometry |  |  |  |  |  |
| RMSD - bonds (Å) | 0.008 | 0.007 | 0.009 | 0.009 | 0.007 |
| RMSD - angles (°) | 0.89 | 0.81 | 0.87 | 0.95 | 0.83 |
| Ramachandran favored (%)** | 97.47 | 96.54 | 96.04 | 94.10 | 96.54 |
| Ramachandran outliers (%)** | 0.00 | 0.31 | 0.33 | 0.00 | 0.31 |
| Poor rotamers (%)** | 1.52 | 5.63 | 0.44 | 1.85 | 5.64 |
| Clashscore | 4 | 10 | 4 | 4 | 8 |
| B-factors(Å<sup>2</sup>) |  |  |  |  |  |
| Mean B-factor | 57.19 | 79.25 | 73.50 | 57.10 | 52.90 |
| Macromolecules | 60.92 | 79.81 | 49.67 | 60.89 | 43.27 |
| Ligands | 49.62 | 0 | 117.7 | 49.50 | 115.40 |
| Solvent | 61.03 | 78.68 | 53.14 | 60.92 | 20.66 |

## RESULTS

### Characterization of Ptet-PolXs enzymatic activities, kinetics, and fidelity

We tested the activity of several N-terminal truncated versions of *Paramecium* PolXs (PolXa-,PolXb-and PolXd-ΔNter) with a relevant DNA substrate, referred to as DSB-cis (**Figure 2A)**. This substrate mimics the DNA ends formed after IES elimination by Pgm in *P. tetraurelia*, featuring the conserved TA dinucleotide and presenting a DSB, with the template nucleotide *in cis* relative to the DNA undergoing extension (like in the gap-filling step on **Figure 1A)**. Without the 2-bp microhomology between the upstream and downstream DNA, it would resemble a short primer extension scenario, which Ptet-PolXs are indeed capable of doing, although not very efficiently (**Figure S1A)**. All DNA polymerases efficiently discriminate dNTPs versus NTPs, as they do not incorporate any ribonucleotide, whether with only GTP or with a mix of NTPs, at any enzyme concentration. This is consistent with the conservation of Polλ’s steric gate residues YFTGS (**Figure S2)**. Moreover, all Ptet PolXs are as efficient with dGTP as with a dNTP mix, leading to a specificity ratio R=[Intensity of the +1 band with dGTP]/[Intensity of the +1 band with dNTP mix] comprised between 0.95 and 1.00.

**Figure 2.**
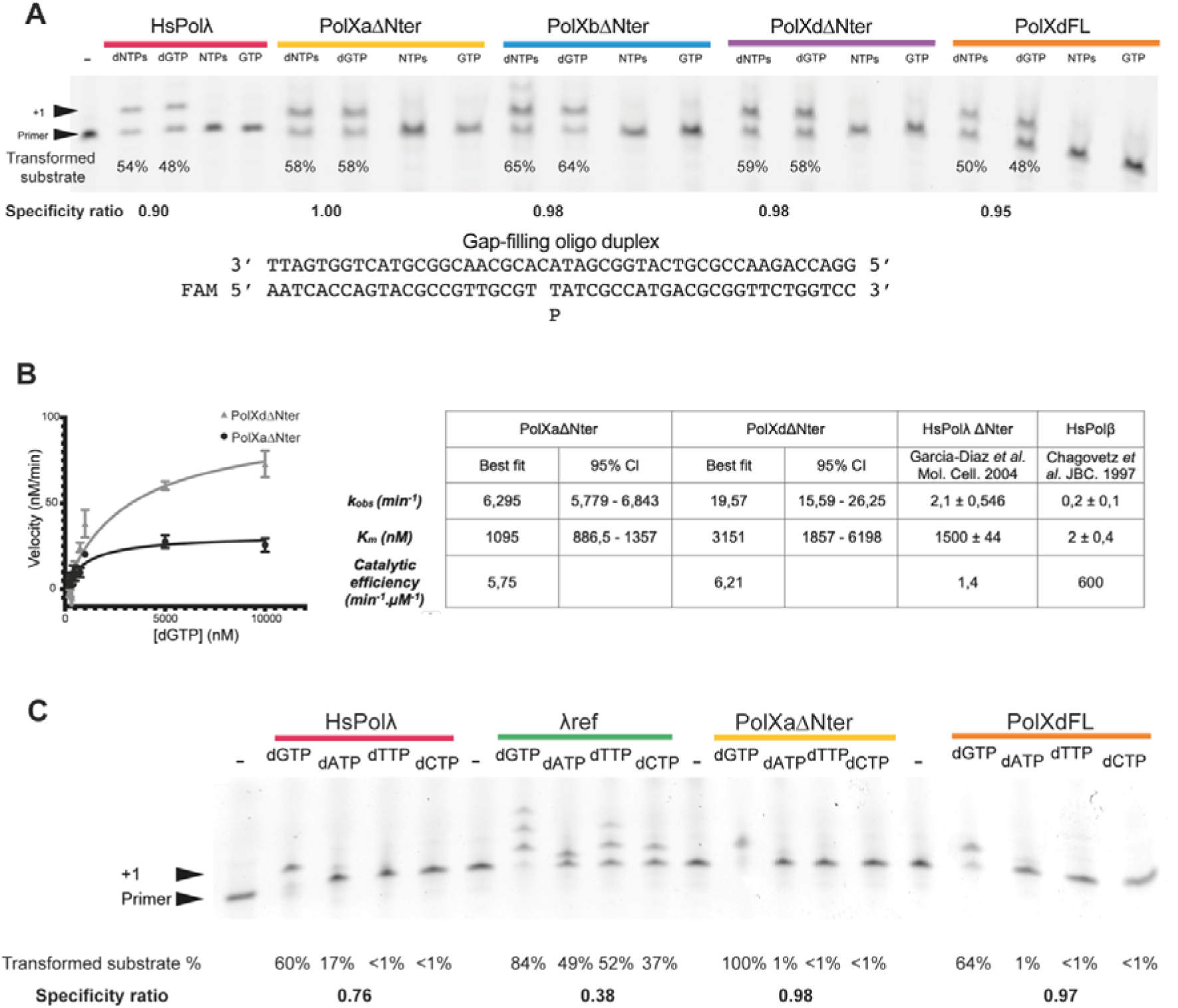
Characterization of Ptet-PolXs enzymatic activities. A: *Paramecium* PolXs display a gap-filling activity similar to human Polλ, independently of their N-terminal BRCT and linker domains. Like HsPolλ, they efficiently discriminate NTPs. Gap-filling assays were conducted using human Pol λ and *Paramecium* PolXaΔNter, PolXbΔNter, PolXdΔNter, and PolXdFL, with either dNTPs, dGTP only, NTPs or GTP only. B: Steady-state kinetics characterization of *Paramecium* PolX. PolXa□Nter and PolXd□Nter have different kinetics profiles in gap-filling, but display a similar catalytic efficiency, which surpass HsPolλ’s. Their behavior in gap-filling is more similar to HsPolλ than to HsPolβ. Activities of PolXaΔNter (n=12) and PolXdΔNter (n=3) were assessed with increasing concentrations of dGTP, employing a gap-filling substrate. Left panel: Velocity curves for PolXa and PolXd, as a function of dGTP concentration. The error bars correspond to the standard deviation on the measurements. Right panel: Kinetic values derived from the plots for each DNA polymerase. The optimal fit values and the 95% confidence intervals (CI) are provided. Literature data for HsPolλ and HsPolβ under analogous conditions are included. C: Fidelity of Ptet-PolXs is higher than HsPolλ’s, independently of its N-terminal BRCT and linker domains. Single time-point, single-turnover fidelity assays were conducted with human Polλ (FL or lacking its N-terminal BRCT and linker domains) and *Paramecium* PolXaΔNter and PolXdFL. The substrate is the gap-filling oligonucleotide duplex indicated in B.

To compare Ptet-PolXs activities with human Polλ, we performed a single nucleotide gap-filling experiment (**Figure 2A)**. In this assay four Ptet-PolXs constructs (PolXa□Nter, PolXb□Nter, PolXd□Nter and PolXdFL) demonstrate comparable activity to HsPolλ. They accurately incorporate a single nucleotide on the FAM-labeled primer with dGTP alone or a mix of dNTPs, and efficiently discriminate NTPs. The specificity ratio is 0.90 for human DNA pol λ.

The N-terminal BRCT and linker domains do not appear to affect catalysis in this context, as evidenced by identical results obtained with both PolXdFL and PolXdΔNter constructs.

We evaluated the dRP lyase activity of the PolXaΔNter construct, considering that its closest metazoan homologues, Polβ and Polλ, both exhibit this activity *in vitro*. In an *in vitro* assay designed to reconstitute a short-range base excision repair context, PolXaΔNter exhibits lyase activity comparable to HsPolβ, indicating a dRP lyase activity (**Figure S1B)**, which is probably shared by the all other Ptet-PolXs, as they also carry the needed residues.

To further compare their activity to human Polλ and β, we characterized the kinetics of incorporation of one representative from each subgroup, PolXaΔNter and PolXdΔNter, under the same conditions as those used previously for human Polλ (53) and Polβ (54), within a gap-filling context. As depicted in **Figure 2B**, PolXaΔNter and PolXdΔNter exhibit distinct catalytic characteristics in this assay: the measured k_obs_ of PolXaΔNter is three times lower than PolXdΔNter but has a three times higher affinity for dGTP, indicated by a higher K_M_. Ultimately, they demonstrate similar catalytic efficiencies of approximately 6 min^-1^.μM^-1^. Comparing these results with previous data reported for human PolX (53, 54), their kinetic parameters more closely resemble those of Polλ than Polβ. Specifically, they exhibit a similar affinity for dGTP as HsPolλ□Nter, with a K_M_ in the millimolar range, but also possess a 3 to 10 times higher turnover number. Consequently, they appear to be 6 times more efficient than HsPolλ□Nter, but they display a low catalytic efficiency compared to HsPolβ.

To assess the fidelity of Ptet-PolXs in comparison to human Polλ, we conducted a qualitative one-point single turnover assay inspired by previous experiments (39). The results obtained for HsPolλ and its mutant λref lacking both N-terminal BRCT and linker domains are in line with previous results from literature (39) (**Figure 2C)**: the full-length enzyme exhibits good fidelity, albeit with a slight dATP incorporation (in line with (55)), while the λref construct appears to make more misincorporations, incorporating several dGTP and dTTP, as well as single dATP and dCTP. In the case of *Paramecium* PolXs, PolXa□Nter and PolXdFL exhibit a common behavior, both of them effectively incorporating the correct nucleotide (dGTP) and not any incorrect dNTP. Thus, Ptet-PolXs demonstrates good fidelity, at least comparable to full-length HsPolλ. Notably, the fact that the fidelity of Ptet-PolXs remains consistent regardless of the presence or absence of the N-terminal BRCT and linker domains, suggests that this fidelity originates only from the enzyme’s polymerase domain.

### Comparison of Ptet-PolXs sequences with other DNA PolX

To understand the position of Ptet-PolXs in the PolX family and the origin of their high fidelity, we used the CLANS classification method (44) on 7250 representative PolXs sequences from diverse organisms (*Paramecium*, metazoan, fungi, viridiplantae, bacteria) obtained by PSI-BLAST. As indicated in **Figure 3A**, 12 clusters were identified manually after convergence of the refinement process using 20000 cycles. We confirmed the accuracy of the method by finding the same relationships between the different subgroups of PolXs that were known and analyzed previously using a smaller dataset (56). **Figure 3A** shows that the metazoan clusters Polμ (#2) and TdT (#1) are close, as expected; the same is true for the Polλ (#5) and Polβ (#8) clusters. Also, other clusters seem to be close to Polλ, such as PolXs from plants (cluster #4), that were classified as Polλ until now (15, 57) or fungal PolXs (cluster #3 and #6) (58). The most divergent PolXs clusters are those including yeast PolIV (or Pol4) and bacterial PolXs. Interestingly, bacterial PolXs are separated in two clusters, containing canonical and non-canonical PolX, as described by Prostova *et al.* (59).

**Figure 3.**
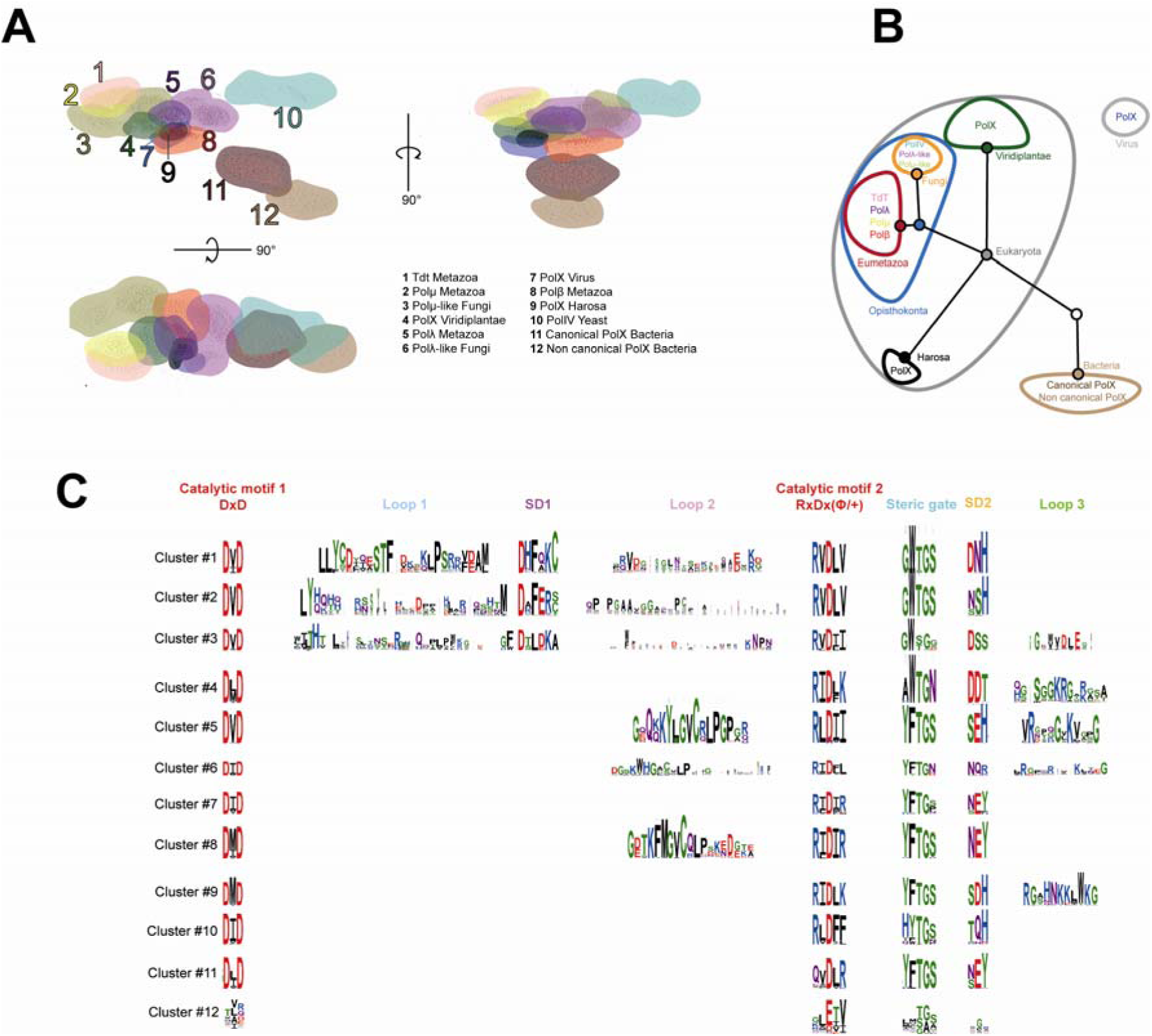
Cluster analysis of PolXs sequences along with the sequence determinants of each subgroup. A: Non-hierarchical clustering of PolXs sequences. A 3D distribution of 7250 PolXs sequences was generated with CLANS (47) using the pairwise Blastp scores of sequence similarity between individual sequences. Three planar projections in different directions of this distribution are shown. The 12 clusters of PolXs sequences are colored and numbered. Their names are given on the bottom right. Ptet-PolXs are in the Harosa group (#9). B: A life tree of the 12 groups of PolXs as determined by the of CLANS. C: Sequence motifs of the 12 PolXs clusters. Each cluster is presented with a sequence logo, where the height of each residue type is proportional to its frequency

We then compared representative sequence motifs of PolXs from each cluster (catalytic motifs, SD1/2 motifs, loops 1/2/3, steric gate motifs). The obtained logos (**Figure 3C)** allowed us to compare the 12 groups and define specific motifs that can be used to determine the group of any new PolX. Those results, along with sequence alignments with known eukaryotic PolXs (**Figure S2)**, indicated that Ptet-PolXs and the other sequences from their group (cluster #9, associated with the clade Harosa that includes ciliates) share similarities with both Polλ and Polβ, including in the steric gate and residues involved in dRP lyase activity. Instead, they appear to be distant from TdT and Polμ as they do not contain Loop1 nor the same SD1 motif (60).

Ptet-PolXs also exhibit a similarity with Polβ in their second catalytic motif (RIDLK). In metazoan Polβ, this motif comprises five conserved residues: RIDIR. The C-terminal arginine residue (R258) plays a crucial role in the induced-fit mechanism of Polβ (30). Unusually among PolXs, *Paramecium* PolXs also display a positively charged residue at this position (K534). They also have an equivalent for all the other residues involved in this induced-fit mechanism: the three catalytic aspartates, the steric gate F506, and a negatively charged residue at the second position of the SD2 motif (NEY in Polβ, SDH in Ptet-PolX). On the basis of this conservation of equivalent residues (especially the positively charged side chain in the second catalytic motif) we formed the hypothesis that Ptet-PolXs could also benefit from an induced-fit mechanism to achieve their high fidelity. In the following, we will test this hypothesis experimentally.

Additionally, Ptet-PolXs display a Loop3 sequence, like metazoan Polλs and related sequences (Plants PolXs, Polλ-like PolXs of Fungi) that is more positively charged than in metazoan Polλ’s (pI of 10.3 vs 8.23 in HsPolλ). Since Loop3 is involved in DNA stabilization during correct catalysis in human Polλ (31), it could have the same role in Ptet-PolX. This hypothesis will also be tested here by site-directed mutagenesis.

### Mutants of Ptet-PolXs and human Polλ designed to probe a Pol β-like induced-fit mechanism

Considering the observed accuracy and fidelity of Ptet-PolXs compared to HsPolλ, we explored the first hypothesis suggested by sequence analysis, involving a polβ-like induced-fit mechanism. First we conducted a fidelity assay to compare the fidelity of WT and mutant versions of PolXa□Nter (**Figure 4A)** involving K534. The PolXaΔNter K534A construct displays strong misincorporations, as it incorporates any dNTP. The mutation at this position also has an effect on the kinetics of the enzyme, since this mutant is the only tested construct to have transformed all of the initial substrates with the correct incoming dNTP. On the other hand, the K534R mutant seems to have a lower effect on fidelity in this assay, and no effect on correct catalysis.

**Figure 4.**
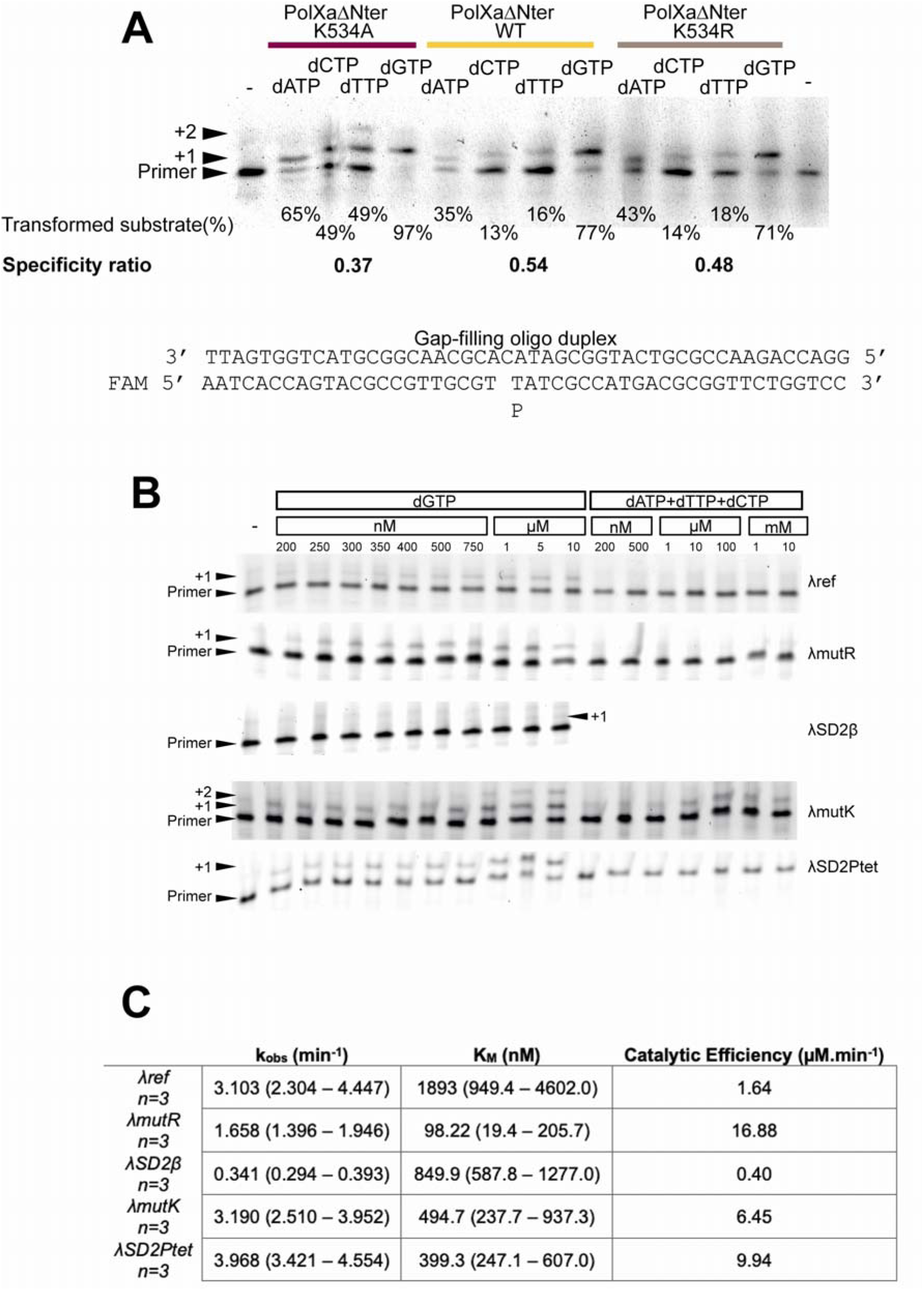
A possible induced-fit mechanism for the fidelity of Ptet-PolXs carried out in part by K534. A: Role of K534 in fidelity for Ptet-PolX. Single time-point, single-turnover fidelity assay conducted with WT or mutant versions of *Paramecium* PolXaΔBRCT. The K534A construct displays a strong error prone behavior, but the K534R mutation has a lower effect on fidelity. B: Gap-filling assays were performed with wild-type (λmut) and different mutant versions of Polλ, with increasing concentrations of correct and incorrect nucleotides. C: Kinetic constants of dGTP insertion by the different mutant versions of Polλ. The value in parentheses stands for standard deviation estimated from n=3.

To further understand this mechanism, we engineered and purified five other mutants of human DNA polymerase λ, the closest relative of Ptet-PolX, at positions believed to confer partly or totally the putative induced-fit mechanism of Polβ. The λmutR construct I492R involves the position equivalent to Polβ’s R258 residue, and the λSD2β mutant has an additional mutation giving it the SD2 motif of Polβ (NEY).

Two other mutants were prepared to give Polλ the equivalent residues of Ptet-PolXs : λmutK has the mutation I492K, and the λSD2Ptet mutant has a second mutation E529D, giving it the SD2 motif of *Paramecium* PolXs (SDH). See Table 1 for a full description of these mutants and their rationale.

After purification, all mutants underwent a steady-state characterization in gap-filling context (**Figure 4B and 4C)** and x-ray crystallography analysis (**Figure 5)** in presence of Ca^2+^ ions to prevent catalysis (61). The λmutR mutant exhibits diminished enzymatic activity compared to the WT enzyme (λref), indicated by a 2-fold lower k_obs_ along with an increased affinity for incoming nucleotides, denoted by a 19-fold lower K_M_. The structural analysis of this mutant (see **Figure 5**, top left panel) revealed that the side chain of the introduced arginine residue is positioned in proximity to the catalytic residue D490 (∼ 3 Å), which is diverted away from the active site. When this mutant is further engineered to include the SD2 motif of Polβ (NEY), it triggers significant differences in enzymatic behavior and local structural changes, but no large movements of domains. This λSD2β mutant displays near-complete inactivity, as evidenced by a 10-fold reduced k_obs_, despite maintaining an almost normal K_M_ in the millimolar range. Crystallographic data for this mutant produced different results compared to other mutants. While all other mutants crystallized in the P2_1_2_1_2_1_ space group with one molecule per asymmetric unit, this specific mutant crystallized in the H32 space group with two molecules in the asymmetric unit, displaying highly anisotropic diffraction data (**Table 2)**, resulting in two slightly different structures (see **Figure 5**, middle and bottom left panels). However, both molecules exhibit the same structural characteristics: closed conformation, absence of an incoming nucleotide in the active site, with residue R492 positioned between the catalytic D490 residue and the NEY SD2 motif, at approximately 3.5-4 Å from both. This proximity suggests potential stabilization through salt bridges, either with catalytic D490 or with both glutamate and tyrosine of the SD2 motif. Additionally, in this instance, Loop3 does not engage with an interaction with the DNA template strand, which is improperly placed for catalysis in the active site (**Figure 6B**).

**Figure 5.**
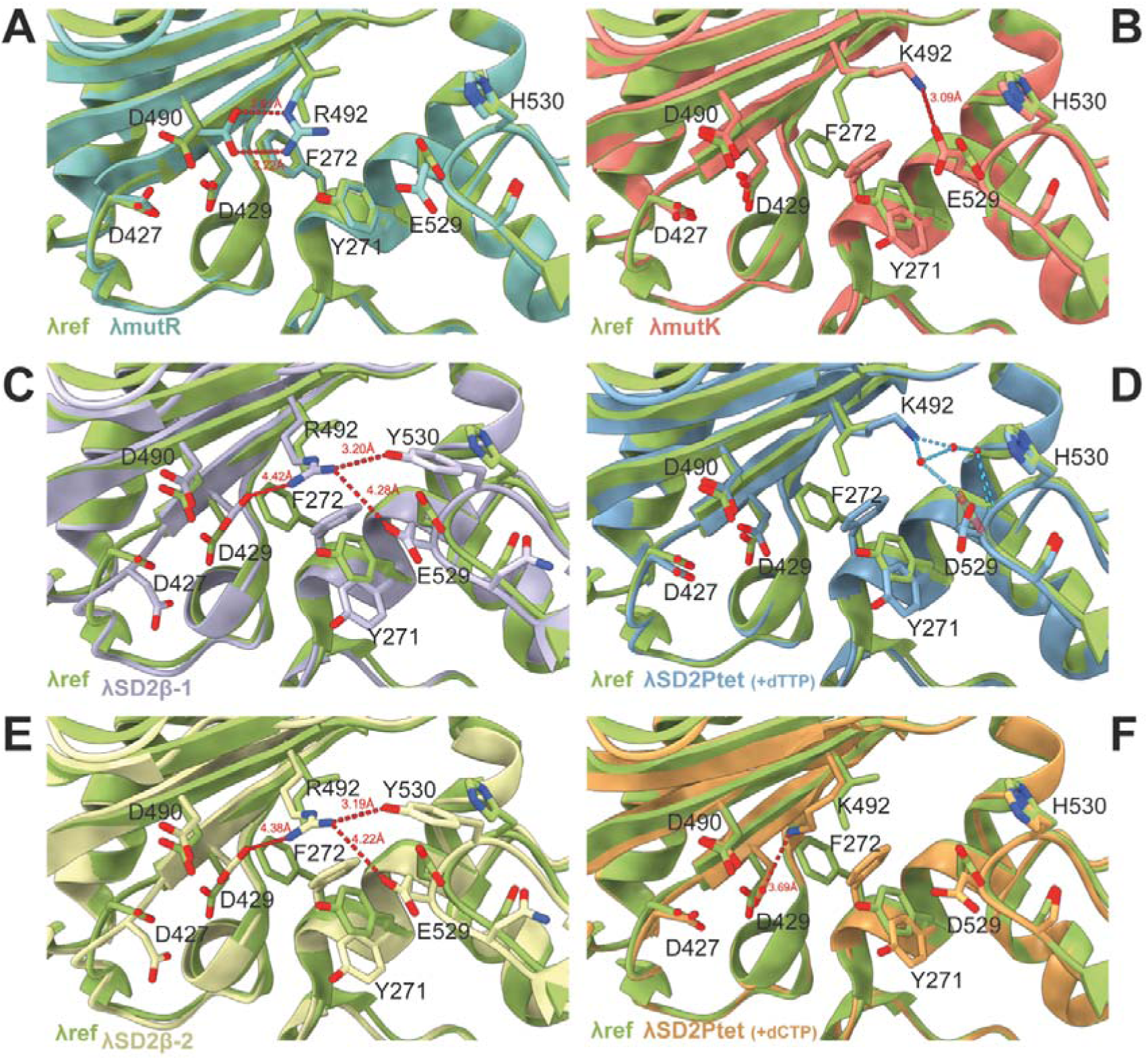
Comparison of the structures of the active site of four mutant versions of Polλ and comparison with λref (PDB 7m43, in green). The residues involved in the activation mechanism of the active site are highlighted, including catalytic residues (D490, D427 and D429), steric gate residues (Y271 and F272), SD2 motif residues (529 and 530), and residue R492 in catalytic motif 2. Distances between residues are indicated in red, while hydrogen bonds are shown in cyan. In the λmutR mutant (top left panel), R492 is oriented towards the catalytic residues at ∼3 Å from D490, which is diverted from the active site. The λSD2β mutant (middle and bottom right panels) displays two molecules in the asymmetric unit. In those structures, D490 is oriented towards R492, which is located between this catalytic aspartate and the SD2 motif, at a distance of ∼3.5-4 Å. In the λmutK construct (top right panel), the inserted lysine (K492) is located at 3 Å from the SD2 residue E529, forming a salt-bridge with it. In the matched (with dTTP) structure of the λSD2Ptet mutant (middle right panel, in blue) K492 is stabilized by the SD2 residue D529 through water-mediated bonds, whereas in the unmatched structure (with dCTP, bottom right panel, in orange) K492 is at 3.7 Å from the catalytic D429, diverting it from the active site.

**Figure 6.**
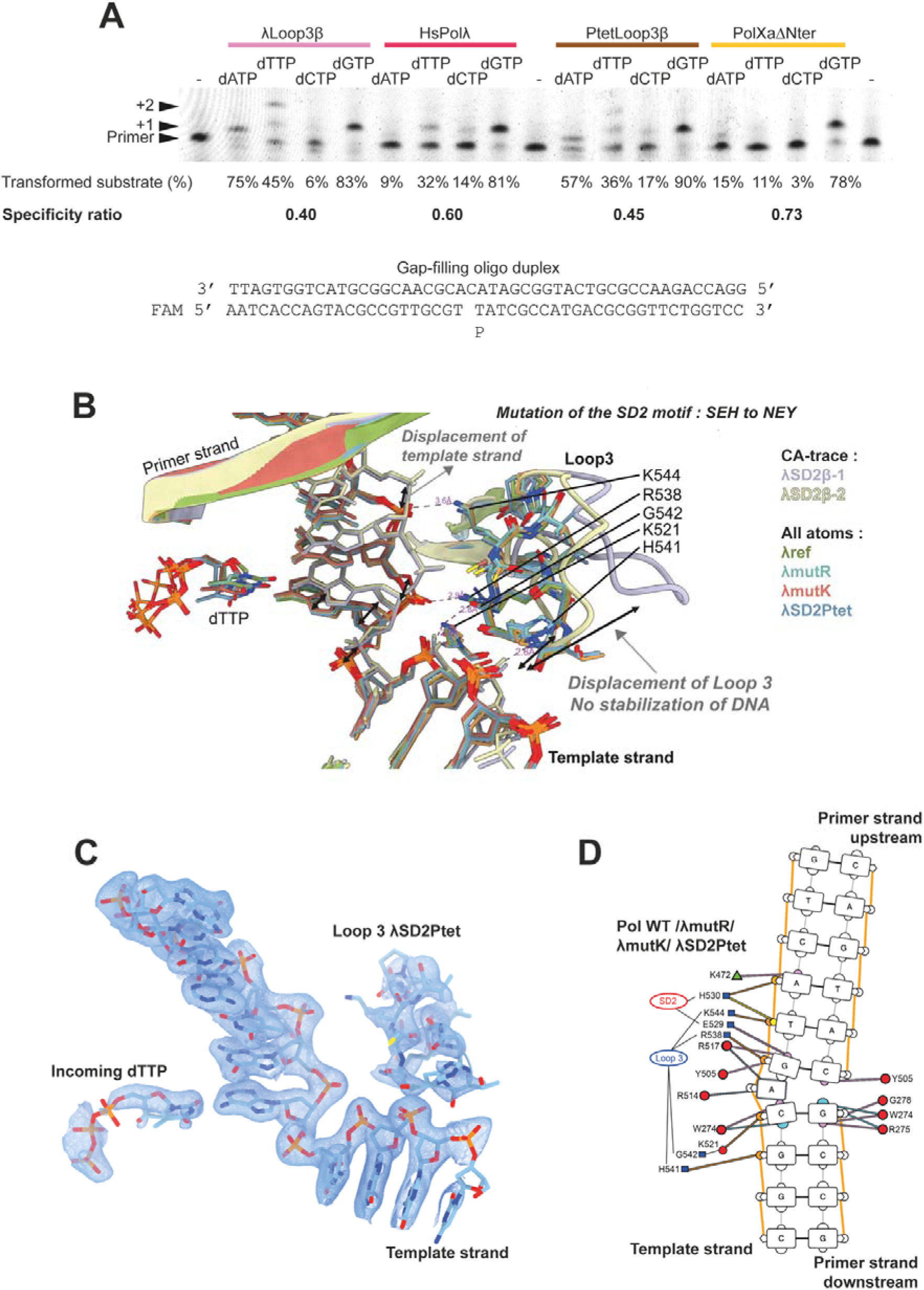
Role of Loop3 in catalysis and fidelity in Ptet-PolXs and human Polλ, and its interaction with the template DNA strand. A: Role of Loop3 in fidelity for Polλ and for *Paramecium* PolX. Top: Single time-point, single-turnover fidelity assays were conducted with human Polλ and *Paramecium* PolXaΔBRCT lacking their respective Loop3 or not. For both enzymes, the truncated constructs display a strong error-prone behavior. Bottom: Sequence of the DNA duplex. B: Comparison of the interaction of Loop3 with the template DNA strand in the six obtained x-ray structures. In λref (PDB 7m43, in green), λmutR (light blue), λmutK (red) and λSD2Ptet (matched in blue, unmatched in orange) structures, Loop3 interacts with DNA. In the λSD2β structure (light purple or yellow), Loop3 is flexible, away from DNA (indicated by bolds arrows) and does not interact with it. Consequently, DNA is displaced and improperly positioned for catalysis (indicated by bolds arrows). C: Example of a 2Fo-Fc electron density map contoured at 1 sigma around the active site, here for the λSD2Ptet mutant. D: The λref, λmutR, λmutK and λSD2β constructs interact with DNA through residues mostly located in the SD2 motif and Loop3. Residues in red circles are located in α helices, those in green triangles are found in β strands, and those in blue squares are found in loops. Interaction of the residues with the DNA double helix are indicated with a color code: interactions in cyan are in the minor groove, those in pink are in the major groove, those in yellow involve the sugar moieties, interactions with bases are in grey, and interactions with phosphate groups are indicated in orange

The λmutK mutant was designed to emulate in Polλ the putative polβ-like induced-fit mechanism of *Paramecium* PolX. Remarkably, its kinetics closely resemble that of the λref construct, particularly in terms of k_obs_, despite exhibiting a 3.8-fold lower K_M_, indicative of a higher affinity for the incoming nucleotide. However, the most notable feature of this mutant is its reduced fidelity: under the same steady-state conditions as those used for testing the correct incorporation of dGTP, it is the only mutant that demonstrated clear misincorporations and double incorporations of the correct nucleotide in the 1-nt gap filling substrate (**Figure 4B)**. Structurally, the mutant closely resembles the WT enzyme with a closed conformation, but the side chain of K492 is participating in a salt bridge with E529 (WT SD2 motif) (see **Figure 5**, top right panel). A significant departure from the WT enzyme is observed in the orientation of the side chains of Y271 and F272. In this mutant (as well as in λSD2β and λSD2Ptet mutants), these two residues adopt a perpendicular conformation resembling that seen in the inactive binary complex of HsPolλ or Polβ with DNA, which is unexpected given the presence of the correct incoming nucleotide.

The λSD2Ptet mutant encompasses all residues potentially implicated in an induced-fit mechanism in *Paramecium* PolX. Its sole sequence difference with the λmutK mutant is the E529D mutation, which mimics the SD2 motif of *Paramecium* PolX. Unlike the λmutK mutant, this variant exhibits no misincorporation behavior and demonstrates kinetics closely resembling λmutK, with slightly improved k_obs_ and marginally lower K_M_, resulting in enhanced catalytic efficiency (1.64 µM^-1^.min^-1^ for λref vs. 9.94 µM^-1^.min^-1^ for the λSD2Ptet mutant and 6.45 µM^-1^.min^-1^ for λmutK). Structural analysis shows two different local conformations within the active site (**Figure S3)**, both present in the structure obtained in presence of the correct incoming nucleotide (dTTP). However, no major conformational changes are observed for this mutant, whose polymerase domain conserves a closed form. In a so called “active” conformation (see **Figure 5**, middle right panel), all catalytic aspartates are correctly oriented in the catalytic site, with K492 engaged in water-mediated bonds with D529 of the SD2 motif. This aspartate participates in bonds not only with its side chain but also with its main chain carboxyl group. In contrast, in an “inactive” conformation (see **Figure 5**, bottom right panel) obtained with an incorrect incoming nucleotide (and within the same structure with the correct nucleotide, see **Figure S3)**, K492 is located 3.7 Å from catalytic D429, which is directed toward this lysine, suggesting the presence of a salt bridge between the two, diverting D429 from the catalytic site. In this scenario, the incorrect nucleotide (dCTP) occupies the active site with an incorrect placement.

In conclusion, these structural and functional studies support a mechanism by which K492 can locally switch between two conformations, an inactive one forming a salt bridge with the catalytic aspartate D429, and a catalytically active one where it is engaged in a completely opposed salt bridge with D529 from SD2 region. This occurs without a big domain rearrangement as seen in polβ.

### Determination of the role of Loop3 in fidelity

To investigate the impact of the conserved Loop3 in the fidelity of Ptet-PolXs and HsPolλ, we generated and purified mutant versions of PolXaΔNter and HsPolλ, lacking their respective Loop3 sequences, namely PtetLoop3β and λLoop3β. We conducted a qualitative fidelity assay to compare the fidelity of the mutant and WT versions (**Figure 6A)** and the results clearly demonstrate that the constructs lacking the Loop3 display a stronger misincorporative behavior. Specifically, the PolXaΔNter construct demonstrates minimal errors, while the PtetLoop3β construct erroneously incorporates dATP, dTTP, and dCTP. A similar trend is observed for HsPolλ, with the mutant construct incorporating dATP and dTTP but seemingly no dCTP. Notably, the correct incorporation of dGTP remains unaffected by the introduced mutations. The predominant misincorporation involves double dTTP incorporation.

Structural analysis allowed us to put forward a tentative explanation for these enzymatic results. For the λSD2β mutant, Loop3 appears to be positioned at a distance from the DNA and exhibits increased flexibility with high B-factors (130 to 260 Å^2^), which contrasts with other structures where it interacts with DNA both upstream and downstream of the templating nucleotide (**Figure 6B)**. Upon Loop3 closure, specific interactions are made with DNA through salt bridges involving its positively charged residues (K544, R538, H541) and DNA phosphates, as well as G542 which interacts with K521 to stabilize a phosphate group (**Figure 6B and 6D)**. Additionally, in those structures, H530 of the SD2 motif also interacts with the phosphate group of the -3 nucleotide (**Figure 6D)**. In the λSD2β structures, which lack this interaction, DNA is displaced, and the templating nucleotide is not properly positioned for catalysis, likely leading to its inability to stabilize an incoming nucleotide.

## DISCUSSION

### *Paramecium* PolXs activity resembles that of human Polλ

We assessed the activity of Ptet-PolXs with their physiological substrate encountered during the programmed genome rearrangement in *P. tetraurelia*. Our results indicate an enzymatic resemblance between these PolXs and both human DNA polymerases β and λ, as they all exhibit gap-filling and dRP lyase activities, and do not incorporate NTPs. Their steady-state characterization, compared with published results obtained for human Polλ (53) and Polβ (54), provide further insights into their enzymatic mechanim and points to a higher similarity to Polλ’s. Thus, it is possible that Ptet-PolXs employ a structural mechanism akin to that of Polλ and other NHEJ-related PolX, maintaining a closed form of their active site throughout catalysis, contrary to what is seen in polβ.

Ptet-PolXs appear to be more accurate than HsPolλ, independently of their BRCT and linker domains.

To understand the origin of this high fidelity, we compared Ptet-PolXs sequences with more than 7000 PolXs sequences from various organisms. The definition of 12 subgroups of PolXs allowed us to study the distinctive features of each group, and the similarities between them. Close examination of the conserved sequences and crucial motifs allowed to make two distinct hypotheses that could explain Ptet-PolXs fidelity: a local induced-fit mechanism, and a conformational change involving Loop3.

### Paramecium PolXs fidelity can be explained by an induced-fit mechanism like in Polβ, but without closing the whole polymerase domain

By testing the effects of point mutations located in the second catalytic motif of Ptet-PolX, we showed that the presence of a positively charged residue at position 5 of the second catalytic motif is necessary for the fidelity of the enzyme. This fidelity mechanism was analyzed by enzyme kinetics and crystallographic structures of mutant constructs of Polλ. Indirect structural evidences, combined with direct enzymatic characterization experiments, allow to suggest that PolXs involved in NHEJ have a local induced fit mechanism, which relies on a lysine residue in the second catalytic mechanism exchanging salt-bridge partner. Importantly, the structural analyses indicated that the steric gate residues were displaced when the residues involved in this mechanism were present : they display an unusual conformation in presence of DNA and of a correct incoming nucleotide, similar to the conformation seen in Polλ in absence of an incoming nucleotide, which allows the fixation of any nucleotide in the active site. In Polλ, such a conformation is associated with a lowering of fidelity, but here it seems that the local induced-fit mechanism can counterbalance this lowering effect.

### Human Polλ and *Paramecium* PolXs may share a crucial conformational change involving Loop3

Fidelity assays on mutants of HsPolλ and PolXa□Nter lacking Loop3 revealed strong error-prone behaviors, indicating a fundamental role of Loop3 in fidelity (**Figure 6)**. We suggest that the mechanism by which this loop stabilizes template DNA could be also dependent on the interaction of DNA with H530, the third residue of the SD2 motif in Polλ and *Paramecium* PolX. Indeed, the interaction of H530 with DNA will be facilitated by the hybridization of the correct incoming nucleotide with the templating nucleotide in the active site. In Polλ lt has been shown that the hybridization of the correct incoming dNTP triggers repositioning of the template DNA (62). This suggests a mechanism whereby, upon the entry of a correct incoming nucleotide into the active site, its hybridization with the templating nucleotide is further stabilized by R556 and R559 (R514 and R517 in Polλ). This repositions DNA and brings it closer to H530, possibly allowing their interaction and triggering a cascading interaction of Loop3 residues with DNA, “closing it” around DNA. Loop3 provides further interactions with DNA, facilitating the catalysis. Previous studies have shown that the role of Polλ’s R517 is to stabilize and control the nascent base pair during dNTP-induced template strand repositioning (62, 63) : when R517 is mutated, DNA does not reposition, neither do the Loop3. This proposed mechanism is in line with the model proposed by Showalter and Tsai (64) : they proposed that the nucleotide binding in the active site of polymerases triggers conformational changes that allow for closing of the active site around substrates, and that fidelity is dictated by the approach of the chemistry step, at which the energy difference between Watson-Crick and mismatched nucleotide incorporations are at the maximum. In Polλ, there is no closing of the polymerase domain; however, the movement of Loop3 mimicks the closing of the tip of the thumb subdomain as in Polβ and has the same consequence: the stabilization of the DNA in the active site (**Figure 7)**.

**Figure 7.**
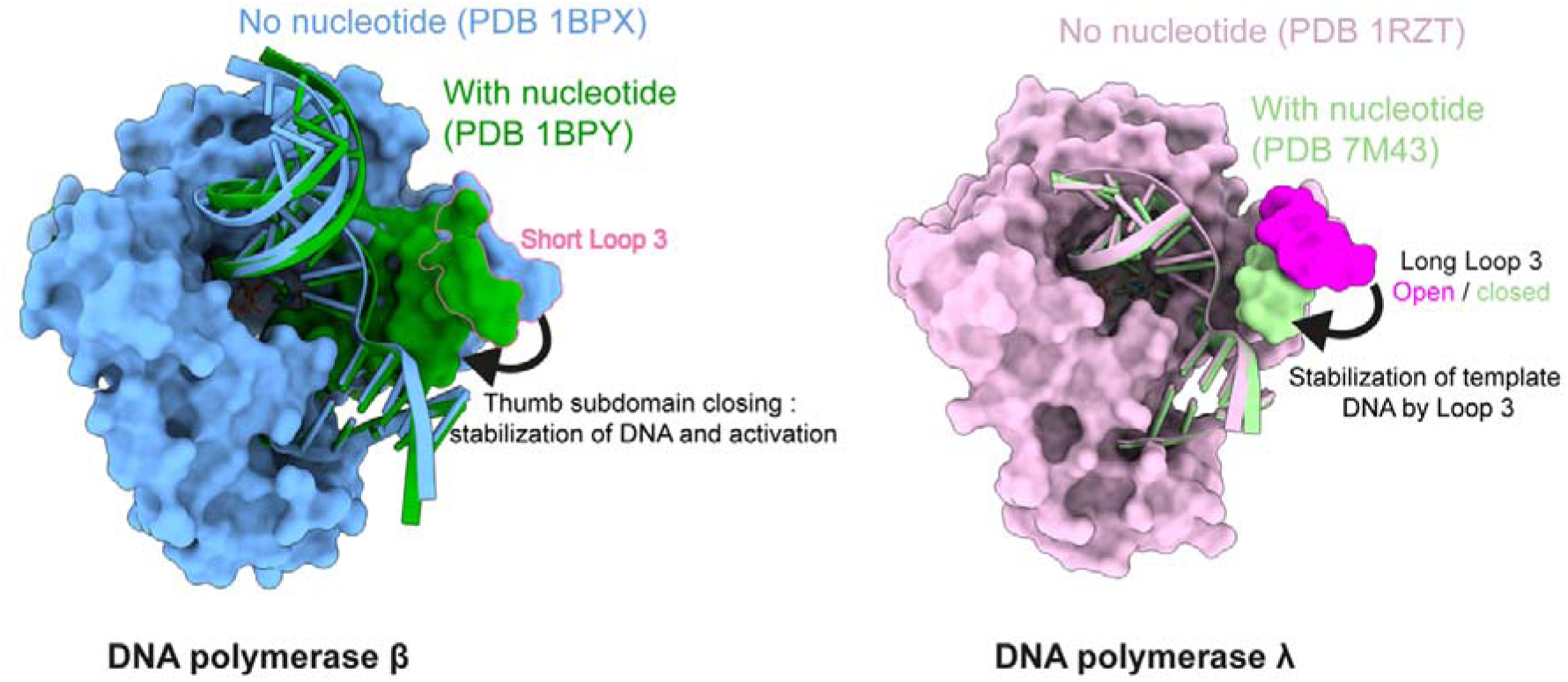
Comparison of conformational changes in the thumb subdomain in Polβ and Polλ. In Polβ (left panel), when the correct incoming nucleotide enters the active site, a global conformational change stabilizes the DNA by a large movement of whole thumb subdomain that closes the polymerase domain. In Polλ (right panel), when the correct incoming nucleotide enters in the active site, only Loop 3 (at the tip of the thumb subdomain) moves in to stabilize the DNA. In this case, the thumb subdomain stays in a closed form.

### A local induced-fit and a conformational change of Loop3 in a single DNA polymerase

The absence of enzymatic activity of the λSD2β construct suggests that the fidelity mechanisms of Polλ (based on Loop3) and Polβ (due to global induced fit and a remodeling of the active site) cannot co-exist in the sequence context of one another, likely because Polλ has evolved towards a permanently closed form and a mechanism relying only on a smaller conformational change. The λSD2β mutant demonstrated an inability to transition to an open state, likely impeded by the permanently closed conformation of Polλ’s catalytic domain (32). As previously discussed, in Ptet-PolXs the induced-fit mechanism is limited to a local side chain movement exchanging salt-bridge partners in the active site, and does not require the opening of the active site to be operative. Instead, the hybridization of a correct nucleotide with DNA can trigger its optimal positioning in the active site, allowing the release of the catalytic aspartate from its interaction with K534. This mechanism, combined with the stabilization of DNA with the Loop 3 in the presence of a correct nucleotide, could explain their high fidelity.

### Conclusion and Perspectives

In summary, our study elucidates that PolXs of *Paramecium tetraurelia* share similarities with both Polλ and Polβ. Notably, they exhibit a fidelity level surpassing that of HsPolλ, a distinctive attribute within the spectrum of NHEJ-associated PolXs. Our studies are compatible with a fidelity enhancement underpinned by two distinct mechanisms, both triggered by the entrance of the correct dNTP and DNA in the active site: a local induced-fit activation of the catalytic site; and a conformational change of Loop 3 originally demonstrated with Polλ by Jamsen *et al*. (31). Even though those conclusions are supported by indirect structural evidence, functional results clearly indicate roles for Loop 3 and residues involved in the local induced fit mechanism in fidelity in Ptet-PolXs. Surprisingly, there are other DNA polymerases of family X that share the residues at the origin of those two fidelity mechanisms: Plants PolXs (cluster #4 in **Figure 3)** display an equivalent of Polβ’s R258 as a lysine and also contain a Loop 3, like HsPolλ. Their steric gate (AWTGN) and SD2 (DDT) motifs also depart from known sequences, so that further studies could reveal a similar function in highly accurate DNA repair. Some fungal “Polλ/µ-like” (clusters #3 and #6 in **Figure 3)** polymerases also seem to share equivalents of Loop3, but with distant SD2 motifs.

On the evolutionary level and based on the fidelity mechanisms in this family of DNA polymerases. *P. tetraurelia* PolXs could represent a missing link in the evolution between Polβ and Polλ.

The CLANS analysis of the whole PolX family could be used to refine the evolutionary scenario proposed by Bienstock *et al.* (56) who proposed that the PolX family originates from bacterial PolX, followed by yeast PolIV, which then separated in eukaryotes in two groups : Polµ/TdT and Polλ/Polβ. We propose that the Polµ/TdT group could originate from the Polµ-like group of PolX found in Fungi; and that eukaryotic Polλ and Polβ could originate from PolX found in Plants and ciliates like *P. tetraurelia*, as those groups display similarities with both Polλ and Polβ (**Supplementary Figure S3B**).

In addition to specific features in the polymerase domain of *Paramecium* DNA polXs the role and structure of their conserved linker domain is particularly intriguing. This domain could play a significant part in the intricate dynamics of the complete NHEJ complex in *Paramecium*, and it strongly differs from that of human Polλ. Disorder predictions and AlphaFold structure predictions suggest that this domain could contain secondary structure elements like α-helices (**Supplementary Figure S1)**. The presence of a structured linker domain between the polymerase and BRCT domains could significantly aid in maintaining the precise orientation of the NHEJ complex and the DNA, thereby enhancing the efficiency of the repair process in the NHEJ complex. Such a structured linker domain could be essential in *Paramecium* ’s specialized NHEJ system to position uniquely and efficiently the DNA substrate and the attached DNA polymerase during DNA ends processing, which is possible because there is only a single type of DNA substrate to deal with. This is in contrast to metazoan Polλ and Polμ, which must handle a wide variety of DNA ends (e.g., varying microhomology lengths, different orientations of microhomologies, …) resulting from various DNA DSB. For *Paramecium* PolX, the ability to stabilize their unique physiological substrate effectively in conjunction with the NHEJ machinery would be highly advantageous. Further structural characterization of the full NHEJ short-range complex in the presence of polX would be needed to answer this question.

## DATA AVAILABILITY

The crystallographic data have been deposited to the PDB under the codes 9EWB, 9EWC, 9EWD, 9EWE and 9EWG.

## SUPPLEMENTARY DATA

Supplementary data are available online.

## AUTHOR CONTRIBUTIONS

Antonin Nourisson: Conceptualization, Data collection and processing, Analysis, Validation, Visualization, Writing—original draft, review & editing. Ahmed Haouz: Data collection and processing. Sophia Missoury: Conceptualization, Methodology, Writing—Review & Editing. Marc Delarue: Conceptualization, Supervision, Writing – Review & Editing.

## ACKNOWLEDGEMENTS

We thank the Molecular Biophysics and Macromolecular Interactions Platform and the Proteomics Platform at Institut Pasteur for help in characterizing the purified proteins by mass spectrometry. We thank the Crystallogenesis and Crystallography Platform (PFX) of Institut Pasteur for help in crystallization and crystallographic data collection. We acknowledge synchrotron SOLEIL (St Aubin, France) for granting access to beamlines Proxima-1 and Proxima-2A, and Pierre Legrand for helpful assistance during the data collection. We thank Mireille Bétermier and Julien Bischerour for many helpful discussions on the NHEJ system of *P. tetraurelia*.

