## Supplementary material for "Mechanisms Ensuring Fidelity of Family X DNA Polymerases in Programmed DNA rearrangements in Paramecium tetraurelia": Suppl Info

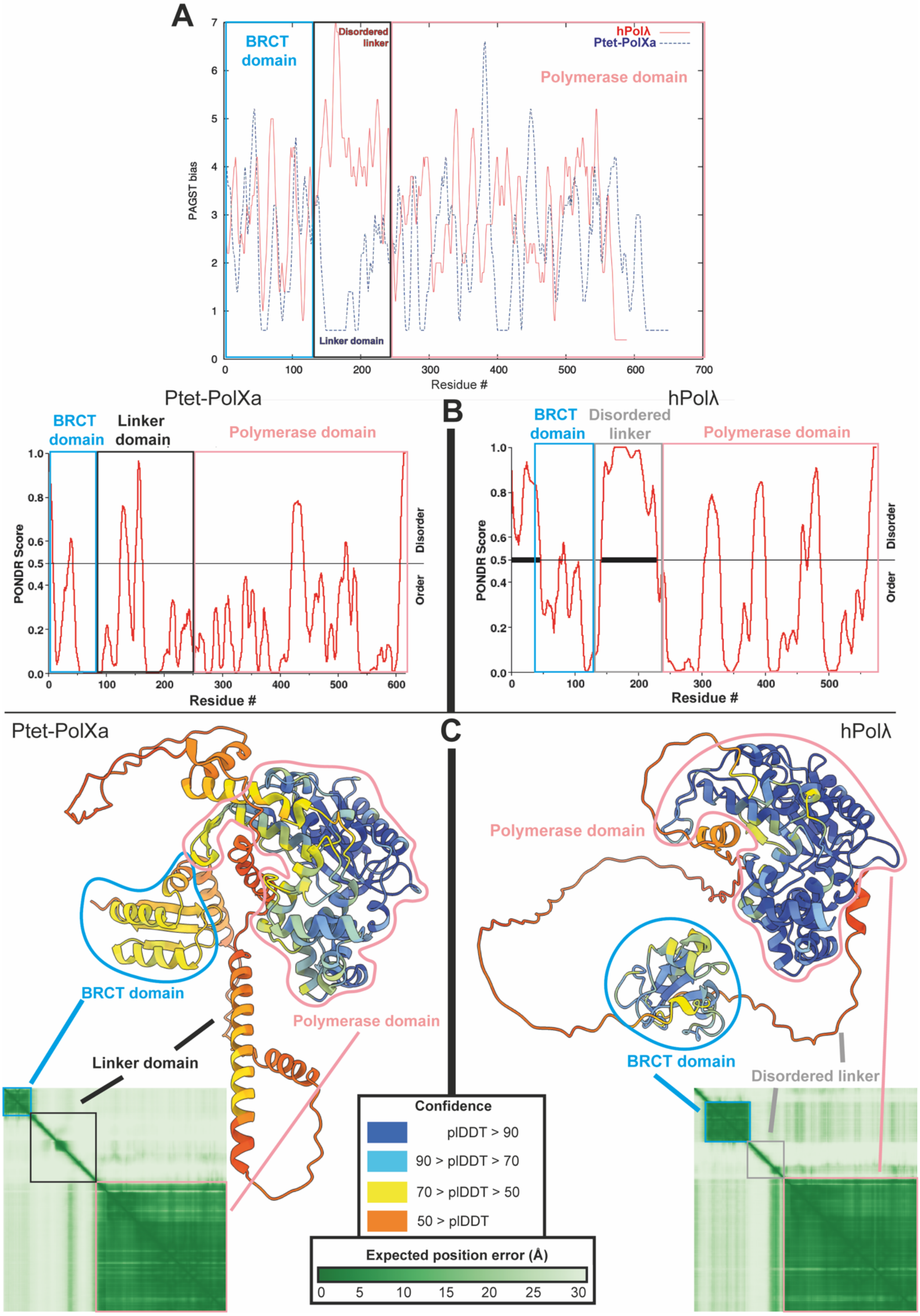

**Supplementary Figure S1. AlphaFold and other predictions of Paramecium PolXs structure.**

**A.** PAGST score of PtetPolx vs human Pol lambda, as a function of residue number. A striking difference is seen in the linker domain, where Pol lambda has an excess of small and polar residues (PAGST) whereas PtetPol have a shortage of such residues. **B**. PONDR (Ref) predicts a clear disordered region in the linker domain for pol lambda, but not for PtetPolX. **C.** AlphaFold3 predicts a number of a-helices in the linker domain of PtetPolX, which anchor relatively clearly the BRCT domain to the polymerase domain, whereas there is no way to predict the orientation of the BRCT domain for human pol lambda, with respect to the polymerase domain.

A

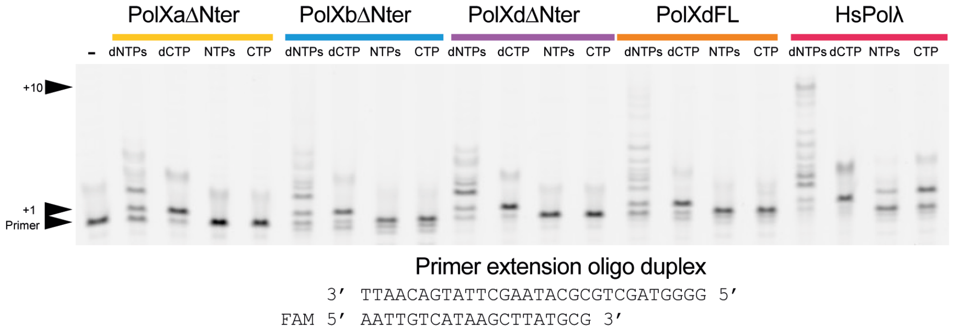


B

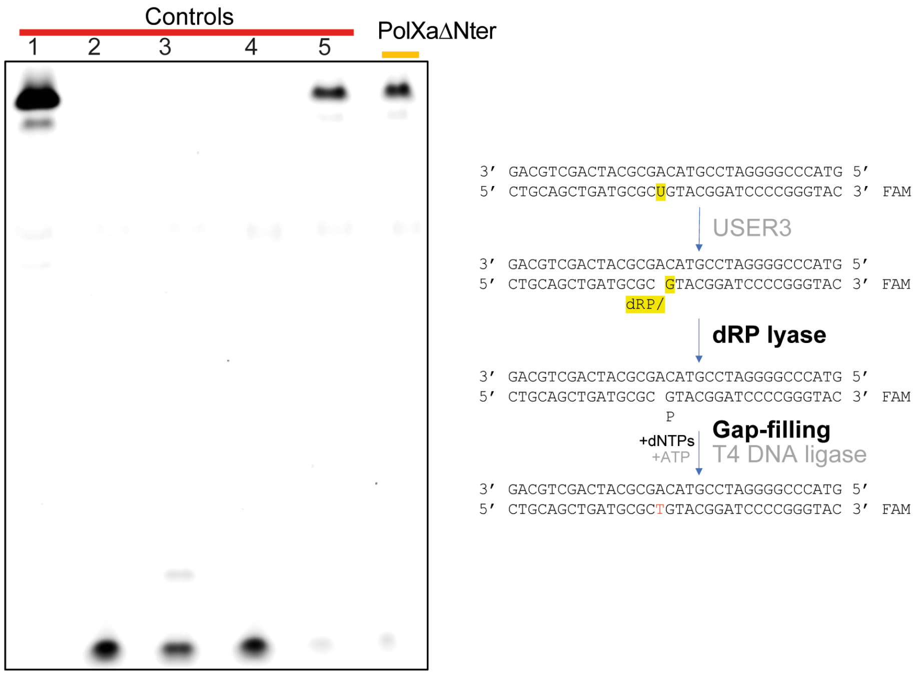


**Figure S2**: Additional enzymatic characterization of *Paramecium* PolX.

**A**: *Paramecium* PolXs can extend a primer DNA strand with dNTPs, albeit inefficiently. Primer extension assay was conducted with 1 µM of either *Paramecium* PolXs and human Polλ in presence of either dNTPs, dCTP only, NTPs or CTP only.

**B**: PolXaΔNter displays a dRP lyase activity like that of human Polβ. The schematic on the right illustrates the principle of this assay. Five control lanes are shown on the gel: 1) DNA prior to USER3 treatment (Full-length with dU); 2) DNA post-USER3 treatment without DNA polymerase or DNA ligase; 3) DNA following USER3 treatment and addition of PolXaΔNter only; 4) DNA following USER3 treatment and addition of T4 DNA ligase only; 5) Positive control featuring human DNA polymerase β.

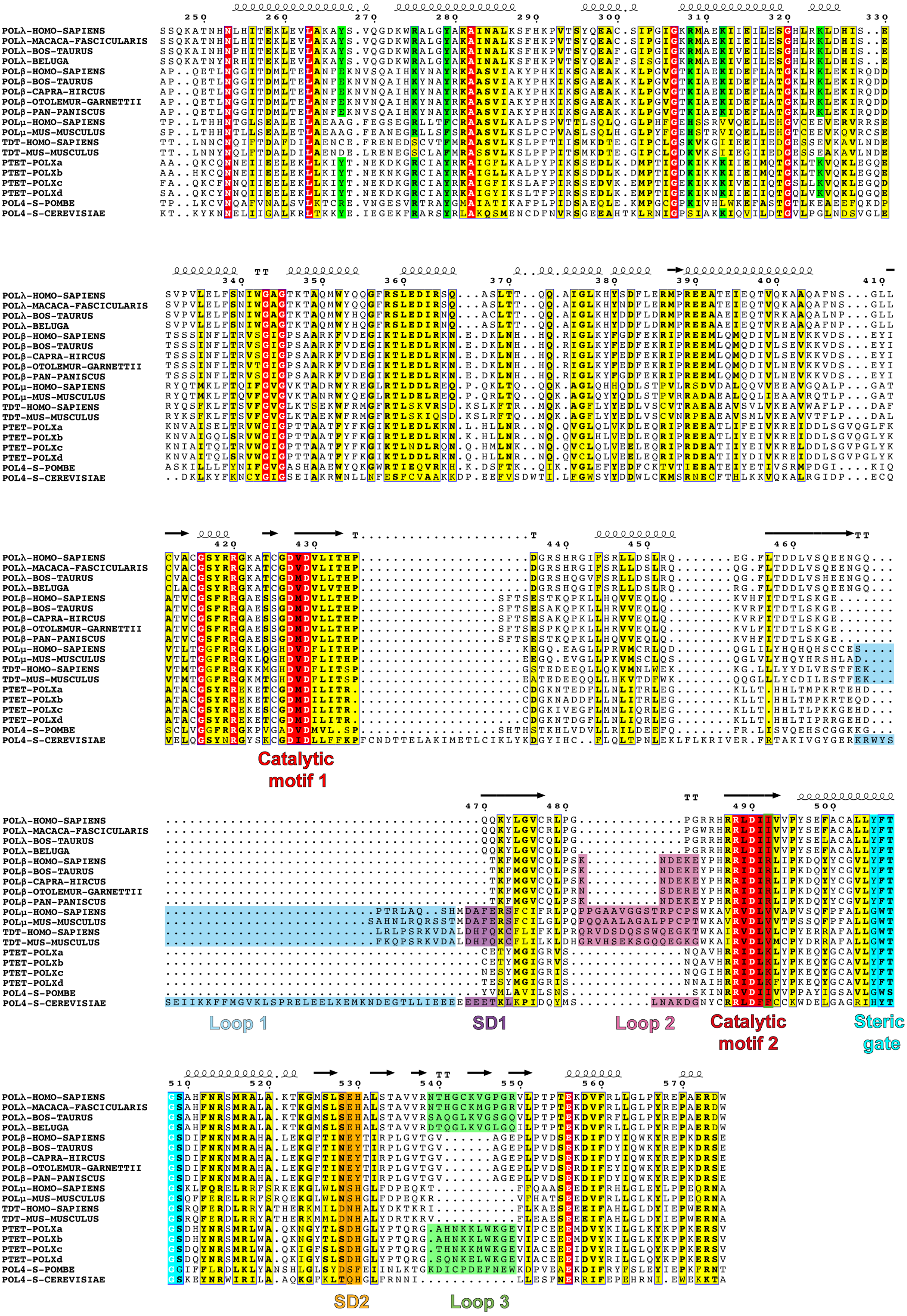


**Figure S3A**: Sequence multialignment of polymerase domains from *Paramecium* PolXs (Ptet-PolXa, b, c, and d) with metazoan DNA polymerase λ, β, μ, and TdT, as well as yeast DNA polymerase IV or Pol4 sequences.

Conserved residues implicated in dRP lyase activity and/or 5'P recognition are highlighted in green. Catalytic motifs DxD and RxDx(φ/+) are highlighted in red. Loop1 is depicted in light blue. The SD1 motif is shown in magenta, Loop 2 is in light pink, and the SD2 is displayed in orange. The steric gate motif is represented in cyan. Loop3 is indicated in light green. Pol4 of *S. pombe* and *S. cerevisiae* have an opposite behaviour in terms of presence/absence of Loop1 or Loop3.

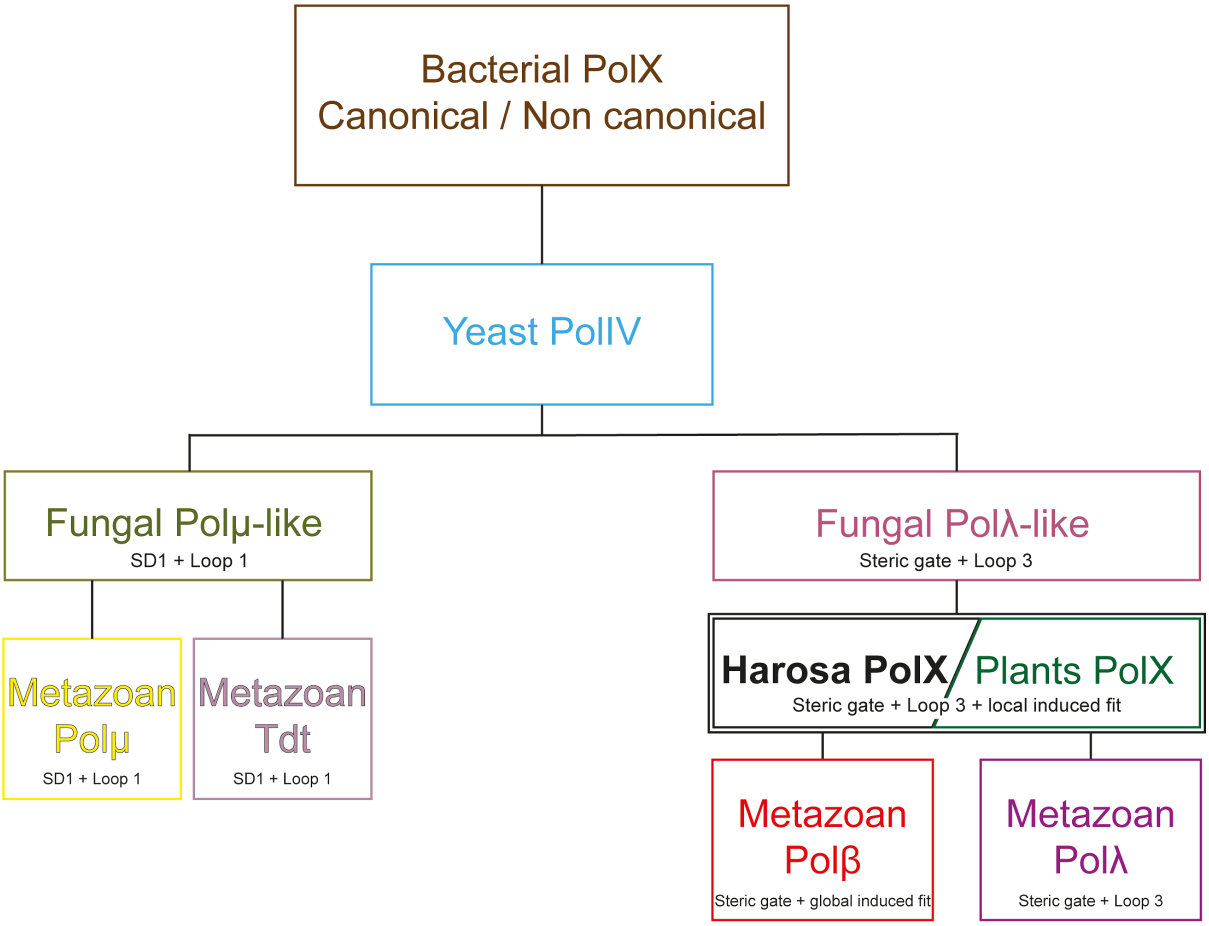


**Figure S3B**. Putative evolutionary tree of all PolXs as inferred from CLANS clusters and the presence/absence of key sequence motifs or characteristic Loops.

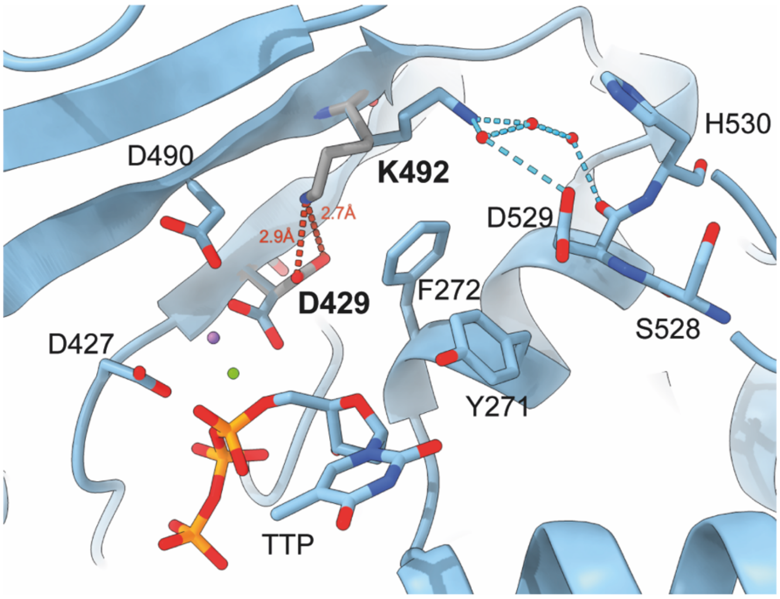


**Figure S4**: The mutant construct λSD2Ptet locally displays two conformations of residues in its active site, in presence of dTTP and DNA. Residues K534 and D429 of its active site display alternative conformations. In the "active" conformation (in blue), K534 is stabilized through water-mediated bonds (light blue) by D529 of the SD2 motif. In the "inactive" conformation (grey), K534 is bonded to catalytic D429 through salt bridges (distances indicated in red), diverting it from the active site. One calcium ion is indicated as a small sphere in purple, and one sodium ion is indicated in green.

**Supplementary Table S1** : Codon optimized nucleotide sequences of the WT constructs used in this study.

| Construct | Nucleotide sequence (codon optimized) |
|---|---|
| PolXaΔNter | AAACAGCAGTTTTGGGAAGCAAAGAAAGGTTATTTTGTTTGTGAAGCCGGTGCAGCACAGAAATGTCAGAATAACGAAATT<br>ATCGAAGAACTGGAAAAGCTGCTGAAGATCTATACCAACGAAAAAGATAAAGGTCGCTGCATTGCATATCGTAAAGCAATT<br>GGTTTTCTGAAAGCACTGCCGTATCCGATTAAAAGTAGCGAAGATCTGAAAGATATGCCGACCATTGGTGACAAAATCAAAA<br>AGAAAATCATCGAAATCATGCAGACCGGCAAACTGACCAAAGTTCAGAAACTGGAAGGTCAAGAGAAAAATGTTGCAATTA<br>GCGAACTGACCCGTGTTTGGGGTATTGGTCCGACCACCGCAGCAACCTTTTATTTCAAAGGTATTAAGACCCTTGAGGACCTG<br>AAAAAGAATCTGCATCTGCTGAATCGTAATCAGCAGGTTGGTCTGCAGCTGGTTAAAGATCTGGAACAGCGTATTCCGCGTG<br>AAGAAGCAACCCTGATTTTTGAAATTGTGAAACGCGAAATCGATGATCTGAGCGGTGTTCAGGGTCTGTTTAAAGCAACCGC<br>ATGTGGTAGCTATCGTCGCGAAAAAGAAACCTGTGGTGATATGGATATTCTGATTACCCGTTGTGATGGCAAAAACACCGAA<br>GATTTTCTGCTGAACCTGATTACACGTCTGGAAGGTAAACTGCTGACCCATCATCTGACCATGCCGAAACGTACCGAACATGA<br>TTGCGAAACCTATATGGGTATTGGCCGTGTTAGCAATCAGGCAGTTCATCGTCGTATTGATCTGAAACTGTATCCGAAAGAAC<br>AGTATGGTTGTGCCGTTCTGTATTTTACCGGTAGCGATCATTATAATCGTAGCATGCGTCTGTGGGCACAGAAAAATGGTTAT<br>ACCCTGAGCGATCATGGTCTGTATCCTACACAGCGTGGTGCACATAACAAAAAGCTGTGGAAAGGTGAAGTTATTCCGTGCG<br>AAGAAGAAATGGACGTCTATAAAATCCTGGGCCTGAAATACAAACCGCCTAAAGAACGTAGCGTGTAA |
| PolXbΔNter | AAACAGCAGTTTTGGGAAGCCAAAAAAGGTTATTTTGTTTGTGAAGCCGGTGCAGCACATAAATGCCAGAATAACGAAATTA<br>TCGAAGAACTGGAAAAGCTGCTGAAGATCTATACCAACGAAAAAAACAAAGGTCGCTGCATTGCATATCGTAAAGCAATTG<br>GTCTGCTGAAAGCACTGCCGTATCCGATTAAAAGCAGTGATGATCTGAAAGATATGCCGACCATTGGCGACAAAATCAAAAA<br>AAAGATCATCGAAATTATGCAGACCGGCAAACTGACCAAAGTTCAGAAACTGGAAGGCCAAGAAAAAAATGTTGCCATTGG<br>TCAGCTGAGCCGTGTTTGGGGTATTGGTCCGACCACCGCAGCAACCTTTTATTTCAAAGGTATTCGTACCCTGGAAGATCTGC<br>GCAAAAACAAACATCTGCTGAATCGTAATCAGCAGGTTGGTCTGCATCTGGTTGAAGATCTGGAACAGCGTATTCCGCGTGA<br>AGAGGCAACCCTGATTTATGAAATTGTGAAACGCGAAATTGATGACCTGAGCGGTGTTCAGGGTCTGTATAAAGCAACCGC<br>ATGTGGTAGCTATCGTCGTGAAAAAGAAACCTGTGGTGATATGGATATTCTGATTACCCGTTGTGATGGCAAAAACGCAGAT<br>GATTTTCTGCTGAACCTGATTACACGTCTGGAAGGTAAACTGCTGACCCATCATCTGACCATGCCGAAACGTACCGAACAGG<br>ATTGCGAAACCTATATGGGTATTGGCCGTGTTAGCAATCAGGCAGTTCATCGTCGTATCGATCTGAAACTGTATCCGAAAGA<br>ACAGTATGGTTGTGCCGTTCTGTATTTTACCGGTAGCGATCATTATAATCGTAGCATGCGTCTGTGGGCACAGAAAAATGGTT<br>ATACCCTGAGCGATCATGGTCTGTATCCTACACAGCGTGGTGCACATAACAAAAAACTGTGGAAAGGTGAAGTTATCCCGTG<br>CGAAGAAGAAATGGACGTTTACAAAATTCTGGGCCTGAAATACAAACCGCCTAAAGAACGTAGCGTGTAA |
| PolXdΔNter | CAGATTCAGTATTGGGAAAACAAACGCGAATTTTTCATTTGTGATGCAGGTAGCGCACAGAAATGCTATAACAACCAGATTA<br>TCGAGGAACTGGAAAAGCTGCTGAAAATCTATACCAACGAGAAAGATAAAGGTCGCTGTATTGCATATCGTAAAGCCATCG<br>GTTATATCAAGAGCCTGACCTTTCCGATTCGTAGCAGCGAAGATCTGAAAGAAATGCCGACCATTGGCGAGAAAATCAAGAA<br>CAAAATCATCGAGATTATCCAGACCGGTCAGCTGGTTAAAGTTCAGAAACTGCAGGGTCAAGAAAAAAATGTTGCAATTACC<br>CAGCTGAGCCGTGTTTGGGGTATTGGTCCGACCACCGCAGCAACCTTTTATTTCAAAGGTATTAAAACCCTGGACGACCTGC<br>GTAAAAATCAGCATCTGCTGAATCGTAATCAGCAGGTTTGTCTGCAGCTGGTTGAAGAACTGGAACAGCGTATTCCGCGTGA<br>TGAAGCAACCATTATCTATGATATTGTGAAACGCGAGATCGATGATCTGAGCGGTGTTCCGGGTCTGTATAAAGCAACCGCA<br>TGTGGTAGCTATCGTCGTGAAAAAGAAACCTGTGGTGATATGGATATTCTGATTACCCGTTGTGATGGCAAAAACACCGATG<br>GTTTTCTGCTGAACCTGATTCAGCGTCTGGAAGGTAAACTGCTGACCCATCATCTGACCATTCCGCGTCGTGGTGAACATGAT<br>ACCGAAAGCTATATGGGTATTGGCCGTATTAGCAATAATGCCATTCATCGTCGTATCGACCTGAAATTCTATCCGAAAGAACA<br>GTATGGTTGTGCCGTTCTGTATTTTACCGGTAGCGATCAGTATAATCGTAGCATGCGTCTGTGGGCACAGAAAATTGGTTATA<br>GCCTGAGCGATCATGGTCTGTATCCGACACAGCGTGGTAGCCAGAATAAAGAACTGTGGAAAGGTGAAGTTATTGCCTGCG<br>AAGAAGAAATTGACGTTTATCGTATTCTGGGCCTGCAGTATAAACCGCCTAAAGAACGTAGCGTTTAA |

| | |
|---|---|
| PolXdFL | ATGTTTAACGCCATCAGCTTTATGTTTCCGCTGAGCACCGTTAATCTGACCAATAATCGTATTAAAACCCTGCGCAATCTGATT<br>GAACGTAATGGTGGTACAATTGAACTGAATAGCCGCACCATTATGATTGTTGGTAGTGATGCAACCAGCGAAAGCTGTCAGA<br>AACAGCTGGAAAAAATGCACCTGAACTTTGAACAGTATCGCCAGCAGTTTATTAACGCAGATTGGATTAGCCAGAGTCTGCA<br>GGCAAAAAATCTGCTGGATTTCAAAAAATACCAGCTGTTCAGCGAAATCGAGCAGAAACAGAAACGTGTTAGTCCGCAGAG<br>TACCGATACCAAATTTGTTTATATTGAGAGCCTGGCAAAAACCGTTCCGCTGAAAGAAGATGCCGAAGATAGCCTGGATATG<br>GAAAGCGGTGAATATACCATTATCAAACCGGAAATGCGCGAGAAATACGAGAAAAAAAAGCAAGAATTCAAACGCCAGATG<br>ATCAAAGAAAATCGTTTCCTGCTGGACTACGAGTATGATAAAGATCTGGATAACTATCACCACCAGGTTGATGATGGTTATG<br>AATTTCTGCTGGACAACTTCCAGATCCTGAAAAAAGAAGAGTTCGACGCACCGCAGCAGAATAAATTCTATGGTGATGAAGA<br>TTGCCAGATCACCGAAATTCGTAAACCGAGCATTGATATTTTTCAAGGTCTGGACATGGGTAAAGCCCTGGTTAATGTTGATC<br>AGCCGCTGGTGAATGCACAGATTAAAGAAACCAAACAGTTTCAGCCTGGTAGCAAACAGCAGATTCAGTATTGGGAAAACA<br>AACGCGAATTTTTCATTTGTGATGCAGGTAGCGCACAGAAATGCTATAACAACCAGATTATCGAGGAACTGGAAAAGCTGCT<br>GAAAATCTATACCAACGAGAAAGATAAAGGTCGCTGTATTGCATATCGTAAAGCCATCGGTTATATCAAGAGCCTGACCTTT<br>CCGATTCGTAGCAGCGAAGATTTAAAAGAAATGCCTACCATCGGCGAGAAGATCAAAAACAAAATCATCGAAATCATCCAGA<br>CCGGTCAGCTGGTTAAAGTTCAGAAACTGCAGGGTCAAGAAAAAAATGTTGCAATTACCCAGCTGAGCCGTGTTTGGGGTAT<br>TGGTCCGACCACAGCAGCAACCTTTTATTTCAAAGGCATTAAGACCCTGGATGACCTGCGTAAAAATCAGCATCTGCTGAATC<br>GTAATCAGCAGGTTTGTCTGCAGCTGGTTGAAGAACTGGAACAGCGTATTCCGCGTGATGAAGCAACCATTATCTATGATAT<br>TGTGAAACGCGAGATCGATGATCTGAGCGGTGTTCCGGGTCTGTATAAAGCAACCGCATGTGGTAGCTATCGTCGTGAAAA<br>AGAAACCTGCGGTGATATGGATATTCTGATTACCCGTTGTGATGGCAAAAATACCGATGGTTTTCTGCTGAACCTGATTCAGC<br>GTCTGGAAGGCAAACTGCTGACCCATCATCTGACCATTCCGCGTCGTGGTGAACATGATACCGAAAGCTATATGGGTATTGG<br>CCGTATTAGCAATAATGCCATTCATCGTCGTATCGACCTGAAATTCTATCCGAAAGAACAGTATGGTTGTGCCGTTCTGTATTT<br>TACCGGTAGCGATCAGTATAATCGTAGCATGCGTCTGTGGGCACAGAAAATTGGTTATAGCCTGAGCGATCATGGTCTGTAT<br>CCGACACAGCGTGGTAGCCAGAATAAAGAACTGTGGAAAGGTGAAGTTATTGCCTGCGAAGAAGAAATTGACGTTTATCGT<br>ATTCTGGGCCTGCAGTATAAACCGCCTAAAGAACGTAGCGTTTAA |
| HsPolλ | GATCCGCGTGGTATTCTGAAAGCATTTCCGAAACGTCAGAAAATTCATGCAGATGCAAGCAGCAAAGTTCTGGCAAAAATTC<br>CGCGTCGCGAAGAAGGTGAAGAGGCCGAAGAATGGCTGAGCAGCCTGCGTGCACATGTTGTTCGTACCGGTATTGGTCGTG<br>CACGTGCCGAACTGTTTGAAAAACAAATTGTTCAGCATGGTGGTCAGCTGTGTCCGGCACAAGGTCCGGGTGTTACCCATAT<br>TGTTGTTGATGAAGGTATGGATTATGAACGTGCACTGCGTCTGCTGCGCCTGCCGCAGTTACCGCCTGGTGCACAGCTGGTT<br>AAAAGCGCCTGGCTGAGCCTGTGTCTGCAAGAACGTCGTCTGGTTGATGTTGCAGGTTTTAGCATTTTTATCCCGAGCCGTTA<br>TCTGGATCATCCGCAGCCGAGCAAAGCAGAACAGGATGCCAGCATTCCTCCGGGTACACATGAAGCACTGCTGCAGACCGC<br>ACTGAGTCCGCCTCCGCCTCCTACACGTCCGGTTAGCCCTCCGCAGAAAGCAAAAGAAGCACCGAATACGCAGGCACAGCCG<br>ATTAGTGATGATGAAGCAAGTGATGGTGAAGAAACCCAGGTTAGCGCAGCCGATCTGGAAGCACTGATTAGCGGTCATTAT<br>CCGACCAGCCTGGAAGGTGATTGTGAACCGAGTCCGGCACCGGCAGTTCTGGATAAATGGGTTTGTGCACAGCCGTCAAGC<br>CAGAAAGCCACCAATCATAATCTGCATATTACCGAGAAACTGGAAGTGCTGGCCAAAGCATATAGCGTTCAGGGTGATAAAT<br>GGCGTGCACTGGGTTATGCAAAAGCAATTAATGCACTGAAAAGCTTCCATAAACCGGTGACCAGCTATCAAGAAGCCTGTAG<br>CATTCCTGGTATTGGTAAACGTATGGCCGAGAAAATCATTGAAATTCTGGAAAGCGGTCATCTGCGTAAACTGGATCATATT<br>AGCGAAAGCGTTCCGGTTCTGGAACTGTTTAGCAATATTTGGGGTGCAGGCACCAAAACCGCACAGATGTGGTATCAGCAG<br>GGTTTTCGTAGTCTGGAAGATATTCGTAGCCAGGCAAGCCTGACCACACAGCAGGCAATTGGTCTGAAACATTATAGCGATT<br>TTCTGGAACGTATGCCTCGTGAAGAAGCAACAGAAATTGAACAGACCGTTCAGAAAGCAGCACAGGCATTTAATAGCGGTCT<br>GCTGTGCGTTGCATGTGGTAGCTATCGTCGTGGTAAAGCAACCTGTGGTGATGTTGATGTTCTGATTACCCATCCTGATGGTC<br>GTAGCCATCGTGGTATTTTTAGCCGTCTGCTGGATAGCCTGCGTCAAGAGGGTTTTCTGACCGATGATCTGGTTAGCCAAGA<br>AGAAAACGGTCAGCAGCAGAAATATCTGGGTGTTTGTCGTCTGCCTGGTCCGGGTCGTCGTCATCGTCGCCTGGATATTATT<br>GTTGTGCCGTATAGCGAATTTGCATGTGCACTGCTGTATTTTACCGGTAGCGCACATTTTAATCGTAGCATGCGTGCCCTGGC<br>CAAAACCAAAGGTATGAGCCTGAGCGAACATGCACTGAGCACCGCAGTTGTGCGTAATACCCATGGTTGTAAAGTTGGTCCT<br>GGTCGTGTTCTGCCGACACCGACCGAAAAAGATGTGTTTCGCCTGCTGGGTCTGCCGTATCGTGAACCGGCAGAACGTGATT<br>GGTAA |

| | |
|---|---|
| HsPolβ | ATGAGCAAACGCAAAGCGCCGCAGGAAACCCTGAACGGCGGCATTACCGATATGCTGACCGAACTGGCGAACTTTGAAAAA<br>AACGTGAGCCAGGCGATTCATAAATATAACGCGTATCGCAAAGCGGCGAGCGTGATTGCGAAATATCCGCATAAAATTAAA<br>AGCGGCGCGGAAGCGAAAAAACTGCCGGGCGTGGGCACCAAAATTGCGGAAAAAATTGATGAATTTCTGGCGACCGGCAA<br>ACTGCGCAAACTGGAAAAAATTCGCCAGGATGATACCAGCAGCAGCATTAACTTTCTGACCCGCGTGAGCGGCATTGGCCCG<br>AGCGCGGCGCGCAAATTTGTGGATGAAGGCATTAAAACCCTGGAAGATCTGCGCAAAAACGAAGATAAACTGAACCATCAT<br>CAGCGCATTGGCCTGAAATATTTTGGCGATTTTGAAAAACGCATTCCGCGCGAAGAAATGCTGCAGATGCAGGATATTGTGC<br>TGAACGAAGTGAAAAAAGTGGATAGCGAATATATTGCGACCGTGTGCGGCAGCTTTCGCCGCGGCGCGGAAAGCAGCGGC<br>GATATGGATGTGCTGCTGACCCATCCGAGCTTTACCAGCGAAAGCACCAAACAGCCGAAACTGCTGCATCAGGTGGTGGAA<br>CAGCTGCAGAAAGTGCATTTTATTACCGATACCCTGAGCAAAGGCGAAACCAAATTTATGGGCGTGTGCCAGCTGCCGAGCA<br>AAAACGATGAAAAAGAATATCCGCATCGCCGCATTGATATTCGCCTGATTCCGAAAGATCAGTATTATTGCGGCGTGCTGTA<br>TTTTACCGGCAGCGATATTTTTAACAAAAACATGCGCGCGCATGCGCTGGAAAAAGGCTTTACCATTAACGAATATACCATTC<br>GCCCGCTGGGCGTGACCGGCGTGGCGGGCGAACCGCTGCCGGTGGATAGCGAAAAAGATATTTTTGATTATATTCAGTGGA<br>AATATCGCGAACCGAAAGATCGCAGCGAATAA |

**Supplementary Table 2** : Constructs and primers used for directed mutagenesis and generation of the mutants constructs of this study. The mutated constructs used for crystallographic structure determination or enzymatic experiments are indicated in **bold**. The other constructs are intermediate constructs used for generation of other mutants.

| Mutated construct | Initial construct | Primers used for mutagenesis |
|---|---|---|
| **PolXaΔNter K534A** | **PolXaΔBRCT** | CGTCGTATTGATCTGGCACTGTATCCGAAAGAACAGTATGG |
| | | ATGAACTGCCTGATTGCTAACACG |
| **PolXaΔNter K534R** | | CGTCGTATTGATCTGAGACTGTATCCGAAAGAACAGTATGG |
| | | ATGAACTGCCTGATTGCTAACACG |
| **PtetLoop3β** | | CGCCACACCACCACGCTGTGTAGGATACAG |
| | | GGTGAAGTTATTCCGTGCGAAGAAGAA |
| PolλΔNter | **HsPolλ** | GCACAGCCGTCAAGCCAG |
| | | CCCTTGAAAATACAGGTTCTCGCCG |
| PolλΔNter-Loop1β | PolλΔNter | GAAACCAAATATCTGGGTGTTTGTCGTCTGC |
| | | GCCTTTAACCAGATCATCGGTCAGAAAACCC |
| **λmut** (ΔNter-Loop1β-C544A) | PolλΔNter-Loop1β | ACAACTGCGGTGCTCAGTG |
| | | GCGTAATACCCATGGTGCGAAAGTTGGTCCTGGTCG |
| **λmutR** (ΔNter-Loop1β-C544A-I492R) | **λmut** | CGTCGCCTGGATATTCGTGTTGTGCCGTATAGC |
| | | ATGACGACGACCCGGACCA |
| **λSD2β** (ΔNter-Loop1β-C544A-I492R-SD2NEY) | **λmutR** | ATGAGCCTGAACGAATATGCACTGAGCACC |
| | | ACCTTTGGTTTTGGCCAGGG |
| **λmutK** (ΔNter-Loop1β-C544A-I492K) | **λmut** | CGTCGCCTGGATATTAAAGTTGTGCCGTATAGC |
| | | ATGACGACGACCCGGACCA |
| **λSD2Ptet** (ΔNter-Loop1β-C544A-I492R-E529D) | **λmutK** | GCCTGAGCGATCATGCACTGA |
| | | TCATACCTTTGGTTTTGGCCAGGG |
| **λLoop3β** (ΔNter-Loop1β-Loop3β) | PolλΔBRCT-Loop1β | GGTCGTGTTCTGCCGACAC |
| | | CGCCACACCACGCACAACTGCGGTGC |
